# Proanthocyanidin-enriched cranberry extract induces resilient bacterial community dynamics in a gnotobiotic mouse model

**DOI:** 10.1101/2021.02.25.432903

**Authors:** Catherine C. Neto, Benedikt M. Mortzfeld, John R. Turbitt, Shakti K. Bhattarai, Vladimir Yeliseyev, Nicholas DiBenedetto, Lynn Bry, Vanni Bucci

**Author notes:** equally contributing authors (Catherine C. Neto, Benedikt M. Mortzfeld). co-corresponding authors; Correspondence should be addressed to: Vanni Bucci, PhD, or, 368 Plantation St Worcester, MA 01605.

## Abstract

Cranberry consumption has numerous health benefits, with experimental reports showing its anti-inflammatory and anti-tumor properties. Importantly, microbiome research has demonstrated that the gastrointestinal bacterial community modulates host immunity, raising the question whether the cranberry-derived effect may be related to its ability to modulate the microbiome. Only a few studies have investigated the effect of cranberry products on the microbiome to date. Especially because cranberry is rich in dietary fibers, we do not know the extent of microbiome modulation that is caused solely by polyphenols, particularly proanthocyanidins (PACs). Since previous work has only focused on the long-term effects of cranberry extracts, in this study we investigated the effect of a water-soluble, polyphenol-rich cranberry juice extract (CJE) on the short-term dynamics of human-derived bacterial community in a gnotobiotic mouse model. CJE characterization revealed a high enrichment in PACs (57% PACs), the highest ever utilized in a microbiome study. In a 37-day experiment with a 10-day CJE intervention and 14-day recovery time, we profiled the microbiota via 16 rDNA sequencing and applied diverse time-series analytics methods to identify individual bacterial responses. We show that daily administration of CJE induces distinct dynamical patterns in bacterial abundances during and after treatment before recovering resiliently to pre-treatment levels. Specifically, we observed an increase of the immunomodulatory mucin degrading *Akkermansia muciniphila* after treatment, suggesting intestinal mucus accumulation due to CJE. Interestingly, this expansion coincided with an increase in the abundance of butyrate-producing Clostridia, a group of microbes known to promote numerous adaptive and innate anti-inflammatory phenotypes.

## Introduction

Cranberry (*Vaccinium macrocarpon*) is a botanical product used worldwide for the maintenance of a healthy urinary tract. It is consumed in the form of fruit, juice, and other products as part of a diet rich in fibers and polyphenols for the prevention of urinary conditions and diseases of aging including cardiovascular diseases and cancers [1]. Cranberry proanthocyanidins (PACs) and other constituents interact with a wide variety of bacteria including gut microbes that cause UTIs and other health conditions, by reducing adhesion, biofilm, co-aggregation [2]. Persistent gut inflammation as experienced in ulcerative colitis, inflammatory bowel disease (IBD) or Crohn’s disease have been linked to genetic factors, lifestyle and dietary habits [3], increasing the risk for colon cancer [4].

Consuming foods high in anti-inflammatory and antioxidant compounds such as polyphenols or dietary fiber may therefore provide a preventative strategy to mitigate these conditions and reduce colon cancer risk. Previous studies by us, using a DSS-AOM mouse model of colitis-induced colon tumorigenesis, showed a significant reduction in colon tumors and tissue inflammation in mice fed either whole cranberry powder[5] or cranberry extracts rich in either polyphenol or non-polyphenol constituents of cranberry[6]. Multiple compounds in cranberries including flavonoids, PACs and triterpenoids have also been reported to reduce tumor cell growth and proliferation, stimulate apoptosis, induce cell cycle arrest and alter associated signaling processes in cells [7–10].

A significant amount of work has recently demonstrated the role of the gastrointestinal microbiota in modulating host immunity[11]. Seminal studies in animal models have demonstrated that short-chain fatty acids, and in particular butyrate-producing Clostridia Cluster IV and XIVa promote the induction of regulatory T-cells and ameliorate symptoms of colitis [12]. Furthermore, these bacteria have been associated with dampening systemic inflammatory response in humans [13] and with promotion of neurological health and of related anti-inflammatory innate immune phenotypes [14]. Recent work has also shown that specific members of the *Bacteroides, Parabacteroides* and *Fusobacterium* genera robustly induce interferon-γ-producing CD8 T cells in the intestine and enhance therapeutic efficacy of immune checkpoint inhibitors in syngeneic tumor models [15]. Similarly, a recent clinical study demonstrated that patients lacking *Akkermansia muciniphila* did not respond to PD-1 checkpoint inhibitor immunotherapy ([16]. Remarkably oral administration *of A. muciniphila* was capable of restoring the efficacy of PD-1 blockade *in vivo* [16], thus demonstrating the causality of the phenotype and highlighting the importance of this bacterium in modulating anti-cancer immunity.

Due to the role that the microbiome has on immune modulation, significant interest is currently placed on understanding the effect of dietary interventions on this system [17] and how diet can be tailored to impact the microbiome and promote health [18, 19]. Thanks to this work it is now established that dietary fibers from plants can promote a healthy and anti-inflammatory microbiome while enrichment in animal diet has been shown to select for bacteria that have been associated with immune dysregulation and pathogenesis [20].

Interestingly, a few studies have investigated the effect of cranberry extracts on the microbiome and shown that members of the genus *Akkermansia*, as well as members of the *Bifidobacteria* and *Clostridia* order appear to be positively affected by long-term interventions with Cranberry derivatives [21–23]. which is also associated with the amelioration of symptoms in a Dextran Sulfate Sodium (DSS)-induced gut inflammation mouse model [24]. However, because cranberry fruit averages about 36% fiber on a dry weight basis [25], we do not know the extent of microbiome modulation that is due to the sole polyphenols. Additionally, it is not known how quickly the microbiome responds to a challenge with polyphenol-rich cranberry extracts, since previous studies only focused on long-term effects. A clearer answer to these questions will provide us with a greater understanding of the role of cranberry polyphenols in modulating gut microbiota dynamics and how cranberry polyphenol-based dietary interventions could be used to promote gut health in the future.

## Results

### Cranberry product composition

A water-soluble, polyphenol-rich cranberry juice extract (CJE) was chosen for this study, allowing for safe administration via oral gavage to gnotobiotic mice. The major polyphenols in cranberries are poly-flavan-3-ol oligomers, or PACs composed primarily of epicatechin units with two types of linkages, either direct carbon-carbon bonding (B-type) or carbon-carbon bonding with an additional ether linkage between units (A-type). Cranberry fruit ranges widely in soluble PAC content depending on cultivar and other factors [26]. PACs are widely distributed in foods and plant sources, and most contain only B-type linkages. The presence of A-type linkages is characteristic of PACs found in cranberries and other *Vaccinium* fruits [27]. PACs have long been associated with the urinary health benefits of cranberry, and cranberry juice and extracts have been the subject of multiple clinical trials and other studies, reviewed in [2]. Constituents detected in utilized CJE are summarized in **Figure 1A**. The total PAC content in the utilized CJE was determined to be 574 ± 40 mg/g (57.4%) using the DMAC method with an authentic cranberry PAC standard. Consistent with previous studies (Patel, 2011)[28], PAC oligomers of up to eight degrees of polymerization with at least one A-type linkage were detected in the 70% acetone-soluble PAC fraction of CJE by MALDI-TOF MS (**Figure 1B,C**) Other polyphenols present in CJE detected by HPLC-DAD and MALDI-TOF MS analyses include flavonols, primarily quercetin glycosides[29] and anthocyanins, primarily cyanidin and peonidin glycosides (**Figure S1, Table S1**). The total flavonol content and total anthocyanin content of CJE were 9.6 ± 0.5 mg/g and 3.4 ± 0.3 mg/g, respectively (**Figure 1A**).

**Figure 1:**
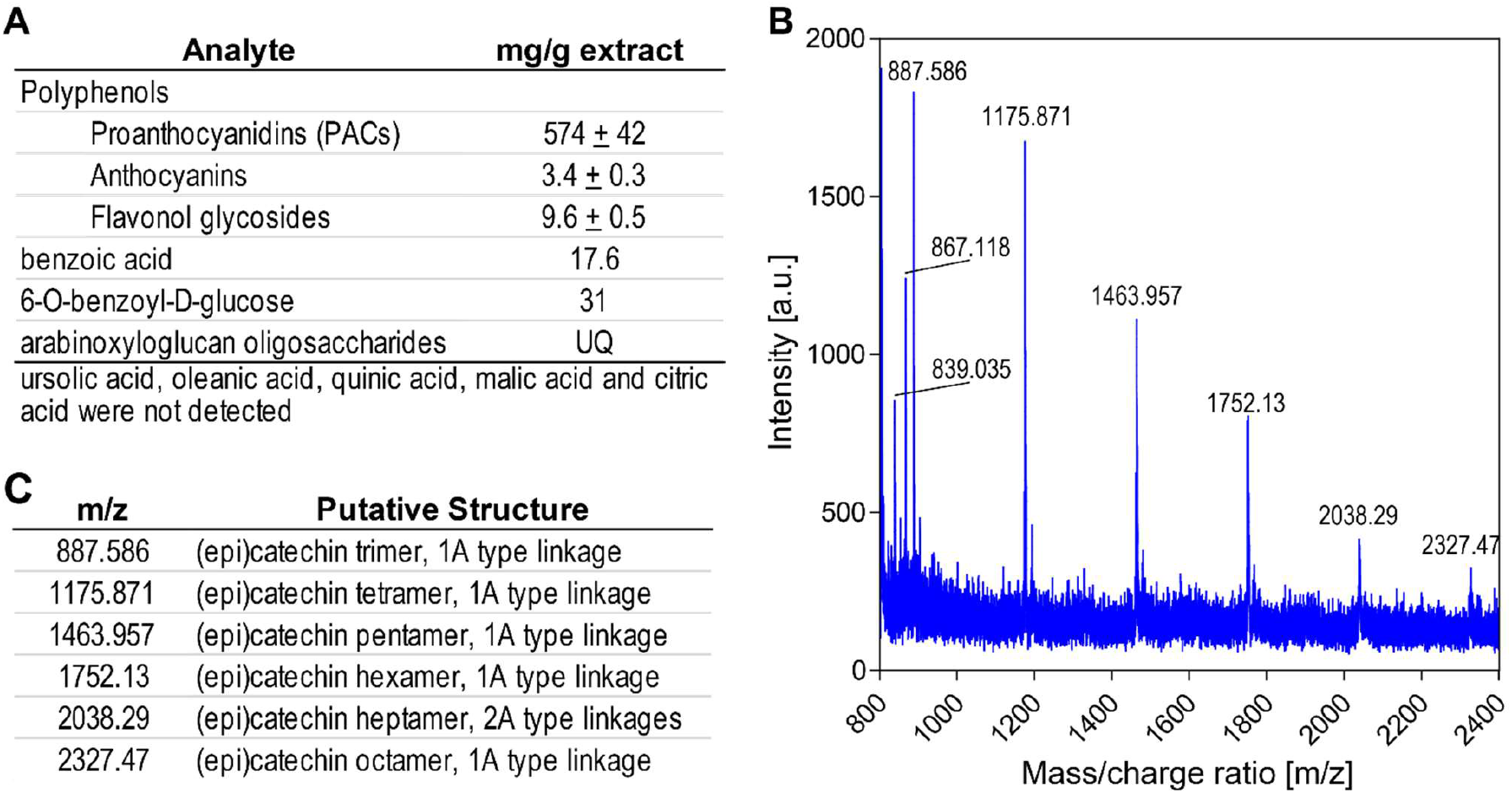
Cranberry juice extract composition. (A) Summary of cranberry juice extract components. UQ=Detected in unknown quantity. (B) MALDI-TOF MS spectrum of proanthocyanidin (PAC) fraction from cranberry juice extract, in positive ion mode. (C) Putative identification of the major ion masses in the PAC fraction.

Quantitative ^1^H NMR analysis found no detectable content of ursolic and oleanolic acids, triterpenoids, which are typically present in the peel of cranberry fruit and associated with the chemopreventive properties of cranberry[6, 7]. Whole cranberry fruit contains approximately 10 mg/g dry weight, or 1% ursolic acid, but due to its low water-solubility, cranberry juice and products derived from juice are much lower in triterpenoid content [8]. No quinic, malic or citric acids were detected, suggesting that smaller organic acids characteristic of cranberry juice were removed by the commercial preparation process (**Figure 1A**). ^1^H NMR confirmed the presence of benzoic acid by comparison with an authentic standard, as well as a major derivative of benzoic acid, the glucoside 6-O-benzoyl-D-glucose, which was identified by comparison of aromatic proton signals between 7.4 – 8.1 ppm and the anomeric proton signal for glucose at 5.6 ppm with those previously reported (**Figure S2**) [30]. The remaining glucoside signals were obscured by other signals in the 3.5 – 5.5 ppm region associated with multiple flavonoid glycosides. Based on peak fit integration of aromatic protons for benzoic acid and its glucoside, the CJE contained 30.9 mg 6-O-benzoyl-D-glucose and 17.6 mg benzoic acid per g dry weight. Thus, CJE contains nearly 5% free and conjugated benzoic acid. ^1^H NMR also contained signals between 6.3 and 6.8 ppm characteristic of p-coumaric acid, a major hydroxycinnamic acid in cranberry, however it appears to be present in very low quantity in CJE.

Multiple ions were detected in the MALDI-TOF MS spectrum of CJE having masses consistent with previously published data for cranberry oligosaccharides (**Figure S3**). These included polygalacturonic acid methyl esters of three and four galacturonic acid units (specifically [M+Na^+^] at 579 for uG3^m2^ m/z = 556; and [M+Na^+^] at 769 for uG4^m3^ m/z = 746) as reported by Sun and coworkers [31] and a series of larger arabinoxyloglucan oligomers containing between 5 – 9 hexose units and 4 – 8 pentose units. This pattern of oligomer masses is similar to those previously reported in cranberry-derived materials [32, 33], but includes larger oligomers, with molecular weights between 1680 and 2532 amu (**Table S2**). Thus, CJE apparently contains a variety of oligosaccharides. We were unable to quantify oligomer content in CJE due to lack of appropriate reference standards.

### Gut microbiome resilience induced by CJE

The current literature reports conflicting results on how cranberry-derived compounds affect the microbial gut community. Most of the microbiome modulatory effect has been attributed to high fiber contents of the fruit as well as their high abundance in polyphenols, however, thorough time-dependent *in vivo* analyses are missing to date, since all previous studies only report analyses through snapshots of selective timepoints [21, 22, 34, 35]. While whole cranberry fruit contains approximately 4% PACs on a dry weight basis [36, 37], other studies have utilized moderately enriched extracts (10 %) in order to investigate the long-term effect of PACs on the microbiome [34] We aimed to study the dynamic response of the gut microbiome to CJE highly enriched in PACs (57 %) over the course of the intervention as well as after the treatment. In order to closely monitor the complex microbial dynamics *in vivo* over several weeks, we chose to utilize a simplified human microbiome consisting of 25 predefined commensal bacteria. Six germ-free C57BL/6J mice were colonized by oral gavage and housed under gnotobiotic conditions. After two weeks of microbiome establishment the mice were given 200 mg/kg body weight (5 mg) of CJE daily for a period of 10 days, followed by a recovery phase of two weeks (**Figure 2A**).

**Figure 2:**
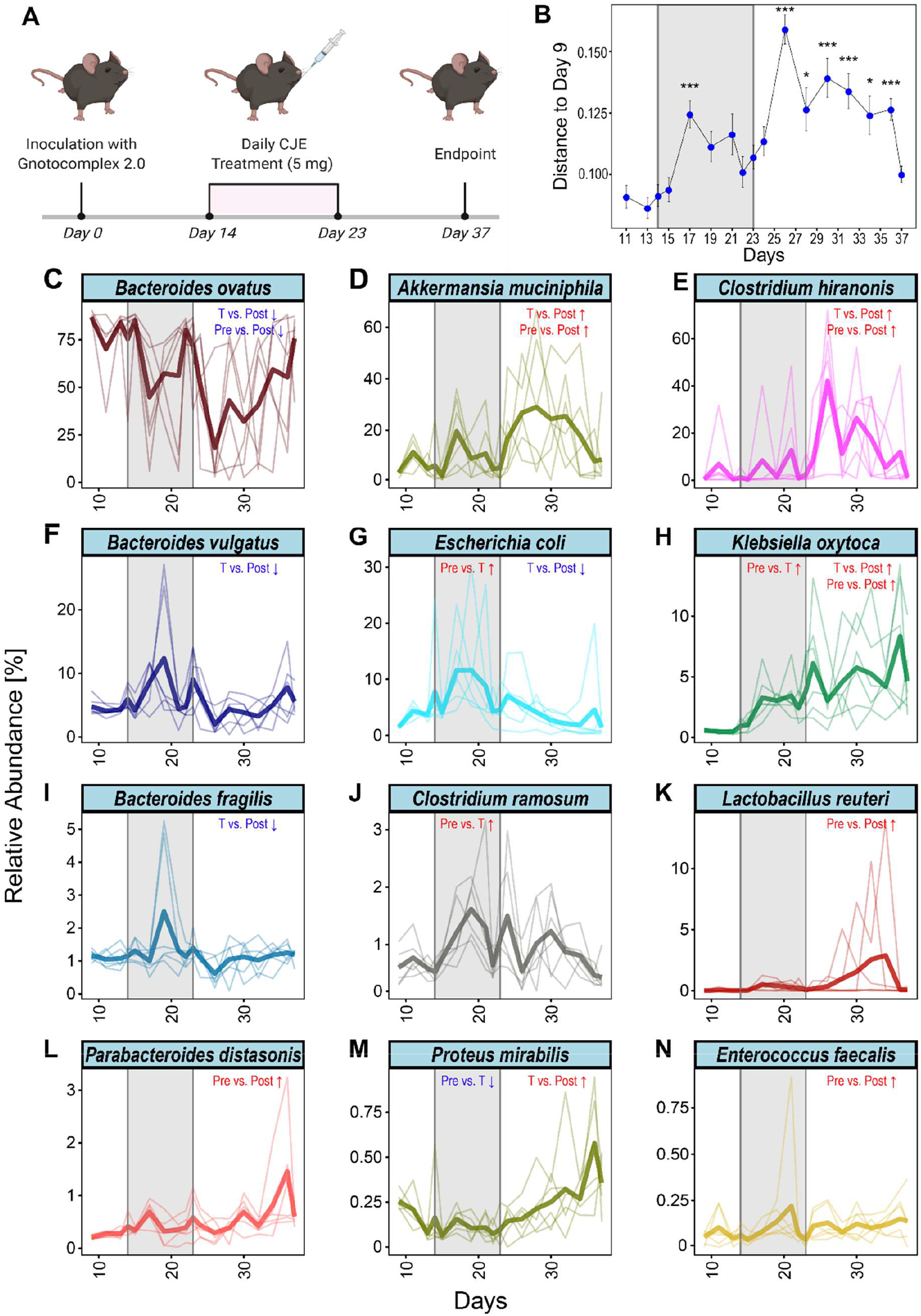
CJE treatment modulates the intestinal microbiome in a gnotobiotic mouse model. (A) Schematic of the experimental setup. (B) Bray Curtis distance of the microbial compositions related to the pre-treatment time point day 9. Points and error bars represent mean and standard error of the mean. *: p≤0.05, ***: p≤0.001. (C-N) Longitudinal depiction of the mean relative abundance throughout the experiment for the 12/15 bacteria that persistently colonized the gnotobiotic mice and were regulated in response to the CJE treatment. Lighter lines show individual replicates. Data for indicated statistical tests are summarized in **Table 1**.

After first establishment of the gut microbiome at day 9 of the experiment, we found that 15 of the 25 species consistently colonized the mice’s guts. Strikingly, the initial microbiome was dominated by *Bacteroides ovatus* with about 80 % total abundance, a prominent colonizer of the human gut microflora. When comparing the Bray Curtis distances of each time point to the pre-treatment on day 9, we found a bimodal dynamic over the course of the experiment (**Figure 2B**). Strikingly, after beginning the treatment with CJE on day 14 we saw a significant increase in the distance to the pre-treatment indicating major changes in the microbial gut composition. Interestingly, the microbiome recovers towards the end of the treatment around day 22 before offset of the CJE intervention induced another jump in distance at day 26. Thereafter the microbiome gradually stabilized resiliently, nearly returning to the pre-treatment level at day 37. In order to statistically evaluate the changes happening throughout the CJE experiment, we leveraged linear mixed effect (LME) modeling to compare the 3 intervals (pre, treatment, post) of the experiment with one another. We found that CJE treatment itself affected species of the family Enterobacteriaceae, as *Escherichia* and *Klebsiella* species are able to significantly increase in abundance together with a gram-positive bacterium *Clostridium ramosum* whereas *Proteus mirabilis* decreases (p<0.05, treatment vs. pre-treatment contrast from LME modeling; see also **Table 1**). As reflected by the distance plot in **Figure 2B**, a greater number of changes was observed when comparing abundances after treatment suspension (after day 23). Specifically, the main colonizer *Bacteroides ovatus* was found to significantly decrease in abundance, coinciding with an increase in *Clostridium hiranonis* and *Akkermansia muciniphila* (p<0.05 post-treatment vs treatment contrast from LME modeling; **Table 1, Figure 2**), making them the most abundant bacteria after *B. ovatus*. While *K. oxytoca* kept increasing in relative abundance even after the treatment, whereas *E. coli* and *P. mirabilis* returned to pre-treatment levels (p<0.05 post-treatment vs treatment contrast from LME modeling). Overall, our data suggest that CJE treatment challenges the dominance of *B. ovatus* and promotes expansion *of A. muciniphila, C. hiranonis* and *K. oxytoca* (p<0.05 post-treatment vs pre-treatment contrast from LME modeling; **Table 1, Figure 2**) in a short time frame in this simplified microbial community.

**Table 1:**
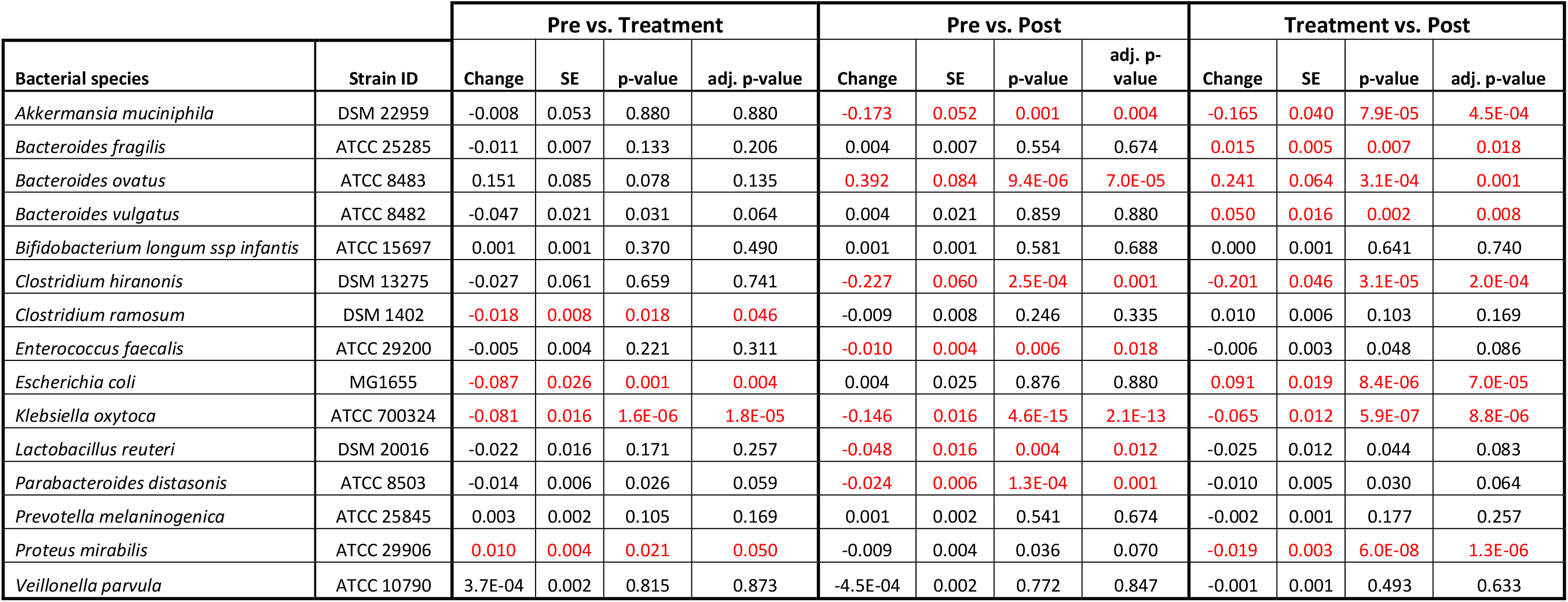
Results of linear mixed effect modeling for the 3 predefined experimental time intervals pre, treatment, post. SE=standard error.

| Bacterial species | Strain ID | Pre vs. Treatment |  |  |  | Pre vs. Post |  |  |  | Treatment vs. Post |  |  |  |
| --- | --- | --- | --- | --- | --- | --- | --- | --- | --- | --- | --- | --- | --- |
|  |  | Change | SE | p-value | adj. p-value | Change | SE | p-value | adj. p-value | Change | SE | p-value | adj. p-value |
| *Akkermansia muciniphila* | DSM 22959 | -0.008 | 0.053 | 0.880 | 0.880 | -0.173 | 0.052 | 0.001 | 0.004 | -0.165 | 0.040 | 7.9E-05 | 4.5E-04 |
| *Bacteroides fragilis* | ATCC 25285 | -0.011 | 0.007 | 0.133 | 0.206 | 0.004 | 0.007 | 0.554 | 0.674 | 0.015 | 0.005 | 0.007 | 0.018 |
| *Bacteroides ovatus* | ATCC 8483 | 0.151 | 0.085 | 0.078 | 0.135 | 0.392 | 0.084 | 9.4E-06 | 7.0E-05 | 0.241 | 0.064 | 3.1E-04 | 0.001 |
| *Bacteroides vulgatus* | ATCC 8482 | -0.047 | 0.021 | 0.031 | 0.064 | 0.004 | 0.021 | 0.859 | 0.880 | 0.050 | 0.016 | 0.002 | 0.008 |
| *Bifidobacterium longum ssp infantis* | ATCC 15697 | 0.001 | 0.001 | 0.370 | 0.490 | 0.001 | 0.001 | 0.581 | 0.688 | 0.000 | 0.001 | 0.641 | 0.740 |
| *Clostridium hiranonis* | DSM 13275 | -0.027 | 0.061 | 0.659 | 0.741 | -0.227 | 0.060 | 2.5E-04 | 0.001 | -0.201 | 0.046 | 3.1E-05 | 2.0E-04 |
| *Clostridium ramosum* | DSM 1402 | -0.018 | 0.008 | 0.018 | 0.046 | -0.009 | 0.008 | 0.246 | 0.335 | 0.010 | 0.006 | 0.103 | 0.169 |
| *Enterococcus faecalis* | ATCC 29200 | -0.005 | 0.004 | 0.221 | 0.311 | -0.010 | 0.004 | 0.006 | 0.018 | -0.006 | 0.003 | 0.048 | 0.086 |
| *Escherichia coli* | MG1655 | -0.087 | 0.026 | 0.001 | 0.004 | 0.004 | 0.025 | 0.876 | 0.880 | 0.091 | 0.019 | 8.4E-06 | 7.0E-05 |
| *Klebsiella oxytoca* | ATCC 700324 | -0.081 | 0.016 | 1.6E-06 | 1.8E-05 | -0.146 | 0.016 | 4.6E-15 | 2.1E-13 | -0.065 | 0.012 | 5.9E-07 | 8.8E-06 |
| *Lactobacillus reuteri* | DSM 20016 | -0.022 | 0.016 | 0.171 | 0.257 | -0.048 | 0.016 | 0.004 | 0.012 | -0.025 | 0.012 | 0.044 | 0.083 |
| *Parabacteroides distasonis* | ATCC 8503 | -0.014 | 0.006 | 0.026 | 0.059 | -0.024 | 0.006 | 1.3E-04 | 0.001 | -0.010 | 0.005 | 0.030 | 0.064 |
| *Prevotella melaninogenica* | ATCC 25845 | 0.003 | 0.002 | 0.105 | 0.169 | 0.001 | 0.002 | 0.541 | 0.674 | -0.002 | 0.001 | 0.177 | 0.257 |
| *Proteus mirabilis* | ATCC 29906 | 0.010 | 0.004 | 0.021 | 0.050 | -0.009 | 0.004 | 0.036 | 0.070 | -0.019 | 0.003 | 6.0E-08 | 1.3E-06 |
| *Veillonella parvula* | ATCC 10790 | 3.7E-04 | 0.002 | 0.815 | 0.873 | -4.5E-04 | 0.002 | 0.772 | 0.847 | -0.001 | 0.001 | 0.493 | 0.633 |

The LME analysis relies on predefined intervals which are set *a priori* to reflect treatment boundaries. However, closer examination of the plots in **Figure 2** reveals that the majority of the observed bacterial dynamics may be shorter than the predefined windows. LME does not find *B. ovatus* to be responding in the during treatment window compared to pretreatment, because its mean relative abundance both drops and recovers throughout the 10 days of CJE intervention.

In order to unbiasedly define intervals of abundance change in the collected time-series, we applied a change point detection algorithm to the data set [38, 39]. Briefly this algorithm infers a position in the time series where the mean of the relative abundance changes across time intervals. For a given number of segments (K), K-1 change points are detected using the dynamic programming algorithm which minimizes the cost of segmentation along with reduced time complexity. To obtain an optimal number of segments (2≤K≤K_max_), an elbow curve is generated using cost of segmentation with respect to the number of segments. A knee point (K_opt_) with maximum curvature is estimated using the maximum of second derivative which is approximated using central difference (**Supplementary Figures S5B to Figure S16B**). Utilizing this approach, we estimated the intervals of change for every bacterium in every mouse (**Supplementary Figures S5 to Figure S16**) as well as for the mean abundance of each bacterium across multiple mice (**Figure 3**). Interestingly, this approach highlights variability across species in their response to CJE in terms of number of occurrences and locations of change points. Investigating the mean change point plots in **Figure 3**, it becomes apparent that three different dynamics can be observed throughout the treatment. Firstly, there was an early response just after the onset of the CJE treatment, followed by a quick partial to full recovery that, in the majority of cases, was still happening during the CJE intervention. This dynamic can be observed for *B. vulgatus, E. coli, B. fragilis, C. ramosum* and *E. faecalis* (**Figure 3, Table 2**). Secondly, we could observe a late response after suspension of the CJE treatment followed by a recovery before the end of the experiment. This dynamic recorded for *A. muciniphila, C. hiranonis* and *L. reuteri* (**Figure 3, Table 2**) may be fueled by the release of the selection pressure imposed by the CJE, allowing for a temporary rearrangement of the microbial community structure post treatment. Lastly, we could observe a set of bacteria that show an early or late response but never experience a recovery in relative abundance, including *B. ovatus, K. oxytoca, P. distasonis* and *P. mirabilis*. Interestingly, even though the change point algorithm does not detect a recovery for *B. ovatus* in the experimental time frame, the data for the individual mice reveal that a recovery event is detected for 4/6 mice before day 35 (**Table 2, Supplementary Figure S7**). Moreover, while the overall pattern looks similar across all mice, the individual responses vary in onset and duration, resulting in a diluted signal and therefore an incomplete recovery in the mean values (**Figure 3A**). Even though the overall resilient bacterial community structure returns back to pre-treatment levels at the end of the experiment (**Figure 2B**), especially the dynamics of the latter bacteria without a recovery demonstrate that an intervention with CJE is able to induce long-term changes in the gut microbiome. Overall, both the treatment with CJE as well as terminating the treatment challenge the dominance of the main colonizer *B. ovatus*, leading to the short-term expansion of other colonizers, including *Bacteroides* species, Clostridia and *Akkermansia*. However, in both instances *B. ovatus* showed signs of recovery within two weeks of change.

**Figure 3:**
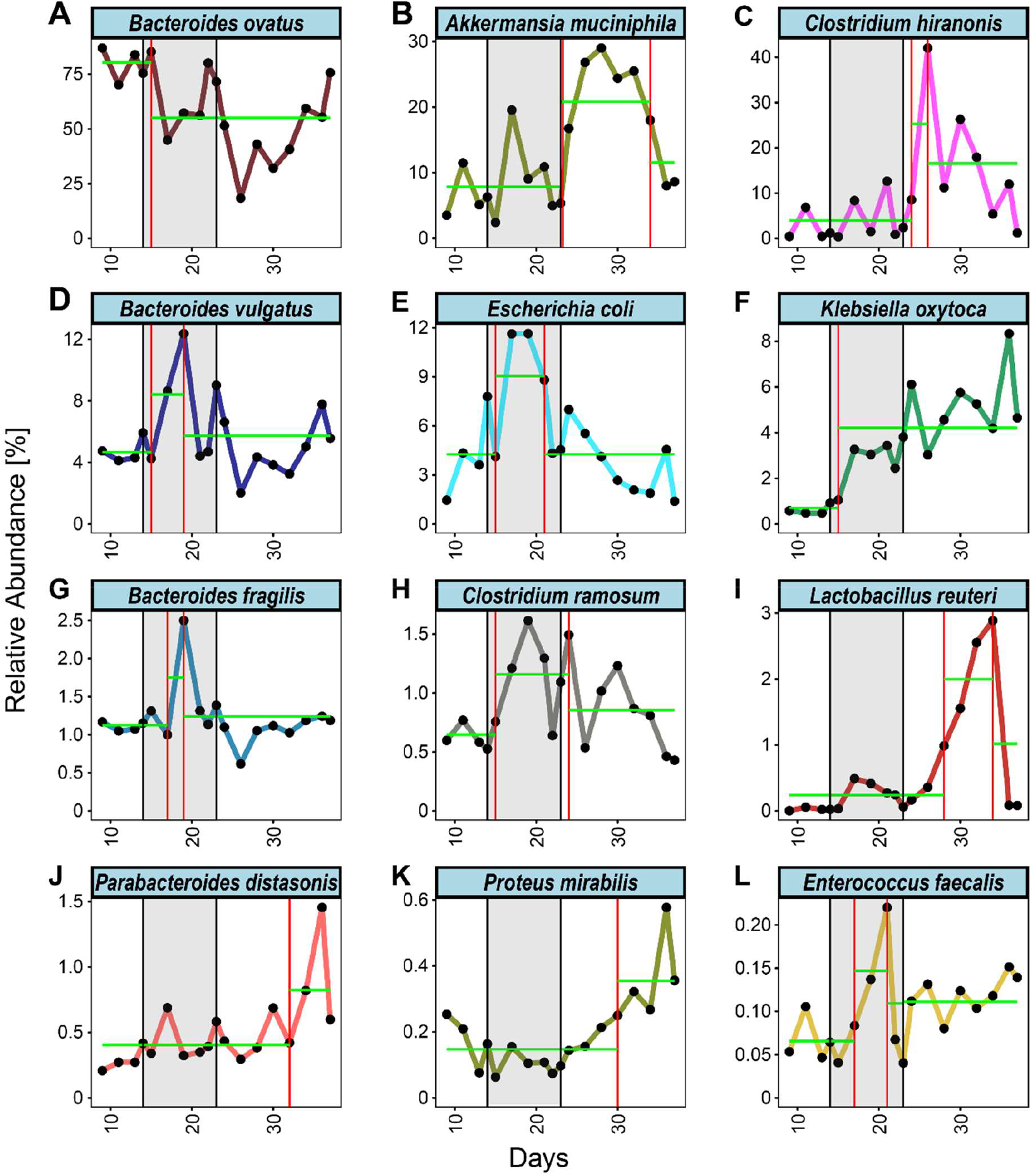
Change point analysis of mean relative abundances throughout the experiment. (A-L) Longitudinal depiction of the mean relative abundance throughout the experiment for the 12/15 bacteria that persistently colonized the gnotobiotic mice and were regulated in response to the CJE treatment. Change points are indicated with a red vertical line, segments are indicated with a green horizontal line. Figures for individual replicates are in the Supplementary Figures, a summary of changes and directions can be found in **Table 2**.

**Table 2:** Results of changepoint analysis describing the dynamics for every bacterium in each mouse. ‘Response’ indicates a change from the pre-treatment level while a ‘recovery’ marks a subsequent change in the opposite direction. Arrow indicates the direction of the response relative to the pre-treatment level.

| Bacterial species | Strain ID | Response/Recovery |  |  |  |  |  | Mean |
| --- | --- | --- | --- | --- | --- | --- | --- | --- |
|  |  | Mouse I | Mouse II | Mouse III | Mouse IV | Mouse V | Mouse VI |  |
| *Akkermansia muciniphila* | DSM 22959 | 2/2 (↑) | 1/1 (↑) | 1/1 (↑) | 1/0 (↑) | 1/1 (↑) | 1/1 (↑) | **1/1 (↑)** |
| *Bacteroides fragilis* | ATCC 25285 | 1/1 (↑) | 1/1 (↓) | 1/1 (↑) | 1/1 (↓) | 1/0 (↑) | 1/1 (↑) | **1/1 (↑)** |
| *Bacteroides ovatus* | ATCC 8483 | 2/2 (↓) | 1/1 (↓) | 1/1 (↓) | 1/0 (↓) | 1/0 (↓) | 2/2 (↓) | **1/0 (↓)** |
| *Bacteroides vulgatus* | ATCC 8482 | 1/1 (↑) | 1/1 (↑) | 1/1 (↑) | 1/1 (↑) | 1/1 (↑) | 1/1 (↑) | **1/1 (↑)** |
| *Clostridium hiranonis* | DSM 13275 | 1/1 (↑) | 1/1 (↑) | 1/1 (↑) | 1/1 (↑) | 1/1 (↑) | 1/1 (↑) | **1/1 (↑)** |
| *Clostridium ramosum* | DSM 1402 | 1/1 (↑) | 1/1 (↑) | 1/1 (↑) | 1/1 (↑) | 1/1 (↑) | 1/1 (↑) | **1/1 (↑)** |
| *Enterococcus faecalis* | ATCC 29200 | 1/1 (↑) | 1/0 (↑) | 1/1 (↑) | 1/0 (↑) | 1/0 (↑) | 1/0 (↑) | **1/1 (↑)** |
| *Escherichia coli* | MG1655 | 1/1 (↑) | 1/1 (↑) | 1/1 (↑) | 1/1 (↑) | 1/1 (↑) | 1/0 (↓) | **1/1 (↑)** |
| *Klebsiella oxytoca* | ATCC 700324 | 1/0 (↑) | 1/1 (↑) | 1/0 (↑) | 1/0 (↑) | 1/1 (↑) | 1/0 (↑) | **1/0 (↑)** |
| *Lactobacillus reuteri* | DSM 20016 | 1/1 (↑) | 1/1 (↑) | 1/1 (↑) | 1/1 (↑) | 1/1 (↑) | 2/2 (↑) | **1/1 (↑)** |
| *Parabacteroides distasonis* | ATCC 8503 | 1/1 (↑) | 1/0 (↑) | 1/0 (↑) | 1/0 (↑) | 1/0 (↑) | 1/0 (↑) | **1/0 (↑)** |
| *Proteus mirabilis* | ATCC 29906 | 1/0 (↑) | 1/0 (↑) | 1/0 (↑) | 1/0 (↑) | 2/2 (↑) | 1/0 (↑) | **1/0 (↑)** |

## Discussion

Cranberry products are consumed around the world for their high nutritional values and antioxidants as well as to prevent urinary tract infections. While it is well established that cranberry derivatives, especially polyphenols, have a modulatory impact on the protective gut microbiome, the mechanisms by which bacteria influence inflammation-linked processes in gut tissues in the presence of cranberry phytochemicals and their various metabolites are not established. Other studies of cranberry’s effect on gut microbiota in various mouse models have reported opposite responses of *Akkermansia muciniphila* in the gut population in response to treatment with cranberry, linking these effects to the polyphenols[34, 40]. However, polyphenol content and composition in cranberry-derived preparations varies widely depending on source materials and method of preparation, but PACs are typically the major constituent by weight[37]. Anhe and coworkers fed C57BL/6J mice on a high-fat high-sucrose diet 200 mg per kg body weight of cranberry extract containing 10% PACs by weight for 8 weeks; the resulting reduction in insulin resistance and intestinal inflammation was associated with a significant increase in *A. muciniphila* ([34]. A related study of cranberry powder in a DSS-treated mouse model of gut inflammation found that the *A. muciniphila* population was boosted significantly by DSS treatment, an effect that could be partially reversed in mice fed cranberry powder for several weeks [40]. While all previous studies focused on long-term microbial effects reporting single time point after several weeks of treatment the short-term effects on the gut microbiome remained unknown. Therefore, we chose a 10-day intervention with a CJE rich in polyphenols (57% PACs) in order to monitor the immediate community dynamics through time and after suspension of the treatment. Using a gnotobiotic mouse model, we did not observe a significant increase of *A. muciniphila* during CJE treatment, however, the bacterium was able to flourish at the expense of the main colonizer *B. ovatus* after the treatment suggesting that it was affected by the PAC-rich CJE during the 10-day intervention.

Cranberry polyphenols have been reported to increase mucin secretion by goblet cells, which helps protect the gut mucous layer and barrier [41]. *Akkermansia* are mucin-degrading bacteria that liberate oligosaccharides from mucin and produce short chain fatty acids [42], which can then be utilized by butyrate-producing bacteria including commensal *Clostridia* (clusters XIVa and IV) and other *Firmicutes* [42]. Interestingly, the expansion of *A. muciniphila* coincides with the expansion of *Clostridium hiranonis* in our study (**Figure 3, Figure S5A Figure S9A**), a cluster XIVa bacterium, whereas *Clostridium ramosum* (Cluster XVIII) spiked during the treatment. Commensal *Clostridia* are strict gram-positive anaerobes that are thought to play important roles in modulating gut homeostasis, maintaining colonocyte health, participation in crosstalk between epithelial and immune cells, and can act as strong inducers of colonic T_regs_ [43]. Low abundance of these *Clostridia* has been linked to inflammatory conditions such as IBD. However, the relationship between *A. muciniphila* and various inflammatory bowel diseases is not completely clear, since overabundance of *Akkermansia* has been reported to exacerbate the inflammation caused by pathogenic bacteria *Salmonella typhimurium* [44, 45].

It is important to note that this and previous studies report relative bacterial abundances without information of actual biomass in the gastrointestinal tract. Therefore, it is possible that certain bacterial species grow in absolute abundance in response to the environmental change, while the main colonizer *B. ovatus* stays unaffected. Nevertheless, it is striking to observe that the mucin-degrading bacterium *A. muciniphila* appears to be kept in check during the CJE treatment even though it has been shown that PAC-related goblet cell density and mucus production in the ileum increase within a few days [41]. This suggests that *A. muciniphila* is susceptible to high concentrations of PACs but can expand in the community after the treatment by degrading the accumulated mucin layer, accompanied by butyrate-producing *Clostridium hiranonis*. However, additional, more detailed longitudinal studies on the impact of cranberry phytochemicals are needed to unravel the mechanisms by which bacteria influence inflammation-linked processes in intestinal tissues and how they manifest in long-term interventions.

In summary our study shows for the first time in a narrow longitudinal data set how a PAC-rich CJE induces community-wide shifts in the intestinal microbiome. Moreover, we are the first to demonstrate that termination of an intervention with a cranberry product induces changes of a magnitude at least as high as the intervention itself. Both intervals (treatment & post) highlight the strong resilience of the gut microbiome which was able to recover close to pre-treatment levels within two weeks. While the dominance of *B. ovatus* is mainly challenged by other *Bacteroides* species, *Clostridium ramosum* and *Escherichia coli* after the onset of the treatment, *Akkermansia muciniphila* and *Clostridium hiranonis* flourish after offset of the selection pressure imposed by the polyphenol-rich cranberry extract.

## Materials and Methods

### Cranberry materials and reagents used in characterization

A food-grade, water-soluble, sterile cranberry-juice derived powder in capsule form (CJE) was donated by Amy Howell of Rutgers University (Ellura®, Trophikos, Inc.). The powder is standardized by the manufacturer to contain at least 36 mg of proanthocyanidins per 240 mg capsule. The capsules were stored at −20°C and in the dark until use. Commercial reagents and standards for analysis were purchased from the following suppliers: Deuterated Dimethylsulfoxide (DMSO-d_6_, 99.9%) and 4,4-dimethyl-4-silapentane-1-sulfonic acid (Cambridge Isotope Laboratories, Andover, MA; N,N-dimethylaminocinnamaldehyde (DMAC), ursolic acid, oleanolic acid (Sigma-Aldrich, St. Louis, MO); malic acid (Eastman Chemicals, Kingsport, TN); citric acid (J.T Baker, Phillipsburg, NJ); quercetin-3-O-galactoside or hyperoside (Chromadex, Irvine, CA); procyanidin-A2 (Indofine Inc., Hillsborough, NJ; quinic acid (Supelco, Bellefonte, PA); cyanidin-3-O-galactoside and peonidin-3-O-galactoside (Extrasynthese, Genay, France).

### Total proanthocyanidin determination

The polyphenol content of the cranberry juice extract (CJE) was determined using established methods. Briefly, total proanthocyanidin (PAC) content was determined using a modification [46] of the industry standard microplate BL-DMAC assay [47]. An isolated whole fruit cranberry PAC fraction prepared as described previously [28] was used as the standard for the DMAC method, and absorbance measurements were obtained using a microplate reader (Molecular Devices SpectraMax M5, SoftMax Pro V5) as described in [36].

### Proanthocyanidin characterization

PACs were isolated from the fraction for further characterization of oligomers by MALDI-TOF MS (Matrix-Assisted Laser Desorption-Ionization – Time-Of-Flight Mass Spectrometry) using methods established previously.[28] Briefly, free sugars were removed from CJE by chromatography on Diaion-HP20, washing with distilled water, then eluting the polyphenols and oligomers using methanol followed by acetone. The eluate was subjected to further chromatography on Sephadex-LH20, eluting with 70:30 methanol/water to remove any residual sugars, phenolic acids and flavonoids, followed by elution of proanthocyanidins using 70:30 acetone/water, rotary evaporation and lyophilization. MALDI-TOF MS analysis was performed by Dr. Stephen Eyles at the University of Massachusetts Amherst Mass Spectrometry Facility using a Bruker Daltonics Omniflex MALDI-TOF mass spectrometer. Data acquisition was carried out in positive ion reflectron mode with 0.1 mM CsI, 0.1% TFA and 50 mM dihydroxybenzoic acid included in the matrix.

### HPLC-DAD analysis

CJE was analyzed for flavonoid composition using HPLC. Identification and quantitation of anthocyanins and flavonol glycosides was performed via reversed-phase HPLC-DAD using a Waters HPLC binary system with 515 pumps coupled with a Waters 996 photodiode array detector and Waters Millenium32 software, as described previously.[36]. Briefly, analyses employed a Waters Atlantis C18 column (100 Å, 3 μm, 3.9 mm x 150 mm) and gradient elution at a flow rate of 0.9 mL/min with mobile phases consisting of 99.5:0.5 (v/v) water:phosphoric acid (A) and 50:48.5:1:0.5 (v/v/v/v) water:acetonitrile:acetic acid:phosphoric acid (B) according to a published gradient scheme as in [48] Flavonol glycosides were detected at a wavelength of 355 nm and quantified based on a quercetin-3-O-glycoside standard; anthocyanins were detected at 520 nm and quantified based on cyanidin-3-O-galactoside and peonidin-3-O-galactoside standards as previously described [6].

### ^1^H NMR analysis

A qualitative profile of CJE was generated, and quantitative NMR to determine several non-polyphenol metabolites was conducted using a Bruker AVANCE III 400 MHz NMR spectrometer equipped with a 5mm BBFO z-gradient probe, as described previously [36]. Briefly, samples were prepared (n=5) at 75 mg/mL in DMSO-d_6_ with 4,4-dimethyl-4-silapentane-1-sulfonic acid as a reference standard. ^1^H NOESY NMR spectra were acquired and processed using TopSpinTM 3.5 and IconNMRTM 5.0.3 as in [36]. Data analysis was performed using AssureNMRTM 2.0 and AMIXTM 3.9.15. Organic acids and triterpenoids were determined by matching signals against a spectral database and quantified using peak fit integration.

### Animal study

To study the dynamics of the microbiota to a polyphenols-rich cranberry extract we adopted an approach similar to that presented in [49]. Briefly, six male germ-free C57BL/6 mice at 8 weeks of age were transferred into individual cages and checked for sterility by plating before the start of the experiment. In order to closely monitor the complex microbial dynamics *in vivo* over several weeks, we chose a simplified human microbiome consisting of 25 predefined species. This allowed us to study the effect of CJE on human gut commensals in an *in vivo* gut environment, simplifying the knowledge transfer to a human study. On day 0 the mice were inoculated with GnotoComplex 2.0 flora by oral gavage [50]. After 14 days, time need to achieve stable bacterial establishment [49], mice were administered daily a dosage of 5 mg (200 mg/kg body weight) via oral gavage (0.250 mL of 20 mg/mL solution) of cranberry juice extract (CJE) for 10 days until day 23 of the experiment. The daily dosage was chosen based on a previously published study in which a similar dosage appears to have been well-tolerated [51]. Fecal samples were collected every two days throughout the course of the experiment and daily around the beginning and and end of the CJE treatment. Fecal pellets were snapfrozen and stored at −80°C until DNA extraction with the DNeasy Powersoil kit by Qiagen (Hilden, Germany) according to the manufacturer’s protocol. Variable region V3 and V4 of the bacterial *16S rRNA* gene were amplified using previously described methods using the universal 341F and 806R primers and sequenced with 300nt paired-end sequences on the Illumina MiSeq platform [52].

### Bioinformatics and computational analyses

Forward and reverse 16S MiSeq-generated amplicon sequencing reads were dereplicated and sequences were inferred using dada2 [53]. Potentially chimeric sequences were removed using consensus-based methods. Resulting amplicon sequencing variants (ASVs) were mapped to the *16S rRNA* gene sequence of the Gnotocomplex 2.0 strains and samples with less than 4000 reads were dropped from the analysis. Sequence files were imported into R and merged with a metadata file into a single Phyloseq object. Due to the repeated-sampling nature of the data (e.g., paired), to determine the effect CJE of each bacterial species abundance we first run linear mixed effect (LME) modeling and predicted the abundance of each bacterium (after applying a square root arcsine transformation) as a function of treatment period (pre/treatment/post) and by using mouse ID as random effect. For each contrast (pre vs. during, pre vs. post, and during vs. post) we used a Benjamini-Hochberg adjusted p-value of <0.05. In addition to running LME which “averages” abundance within a certain time-window we analyzed the abundance data using the change point detections algorithm to detect abrupt shifts in relative abundance of species across different time points (http://sia.webpopix.org/changePoints.html) [38, 39]. Detection of change point is based on the changes in means of relative abundance across time intervals. For a given number of segments (K), K-1 change points are detected using dynamic programming algorithm which minimizes the cost of segmentation along with reduced time complexity. To obtain an optimal number of segments (2≤K≤K_max_), an elbow curve is generated using cost of segmentation with respect to the number of segments. A knee point (K_opt_) with maximum curvature is estimated using the maximum of second derivative which is approximated using central difference.

## Acknowledgements/Funding Sources

The authors wish to acknowledge Amy Howell of Rutgers University for providing the cranberry product, and the support of the Leo and Anne Albert Charitable Trust and the Commonwealth of Massachusetts, Department of Public Health (INTF4005HH2W20081179). Mass spectral data were obtained at the University of Massachusetts Mass Spectrometry Center and samples were sequenced at the Center for Microbiome Research (CMR) at the University of Massachusetts Medical School. Figure 2A was created with BioRender.com.

