## Supplementary Materials for "Proanthocyanidin-enriched cranberry extract induces resilient bacterial community dynamics in a gnotobiotic mouse model"

[*]equally contributing authors

[**]co-corresponding authors

Correspondence should be addressed to:

Vanni Bucci, PhD

**Running title:** Cranberry juice extract microbiome dynamics

**Table S1: Putative identification of ion masses in MALDI-TOF MS spectrum of Figure S1.**

| m/z | Putative Assignment [M+Cs$^+$] |
|---|---|
| 548.986 | Benzoyl-hexoside-pentoside |
| 550.558 | Cyanidin-3-arabinoside |
| 565.050 | Peonidin-3-arabinoside |
| 580.781 | Cyanidin-3-galactoside |
| 596.946 | Peonidin-3-galactoside / Quercetin-hexosides |
| 613.021 | Myricetin-hexoside |
| 653.066 | Unknown |
| 667.079 | Unknown |
| 700.964 | Quercetin-3-O-(6”-benzoyl)-b-galactoside |
| 709.044 | Procyanidin A2 |
| 742.965 | Quercetin-3-O-(6”-p-coumaroyl)-b-galactoside |

**Table S2: Putative identification of ion masses in MALDI-TOF MS spectrum in Figure S3.**

| m/z | Putative Structure [M+Cs$^+$] |
|---|---|
| 996.997 | (epi)catechin trimer |
| 1285.050 | (epi)catechin tetramer |
| 1489.223 | Xyloglucan, $Hex_5Pent_4$ |
| 1813.307 | Xyloglucan, $Hex_7Pent_4$ |
| 1945.337 | Xyloglucan, $Hex_7Pent_5$ |
| 2077.369 | Xyloglucan, $Hex_7Pent_6$ |
| 2209.397 | Xyloglucan, $Hex_7Pent_7$ |
| 2239.400 | Xyloglucan, $Hex_8Pent_6$ |
| 2341.400 | Xyloglucan, $Hex_7Pent_8$ |
| 2371.432 | Xyloglucan, $Hex_8Pent_7$ |
| 2503.455 | Xyloglucan, $Hex_8Pent_8$ |
| 2533.463 | Xyloglucan, $Hex_9Pent_7$ |

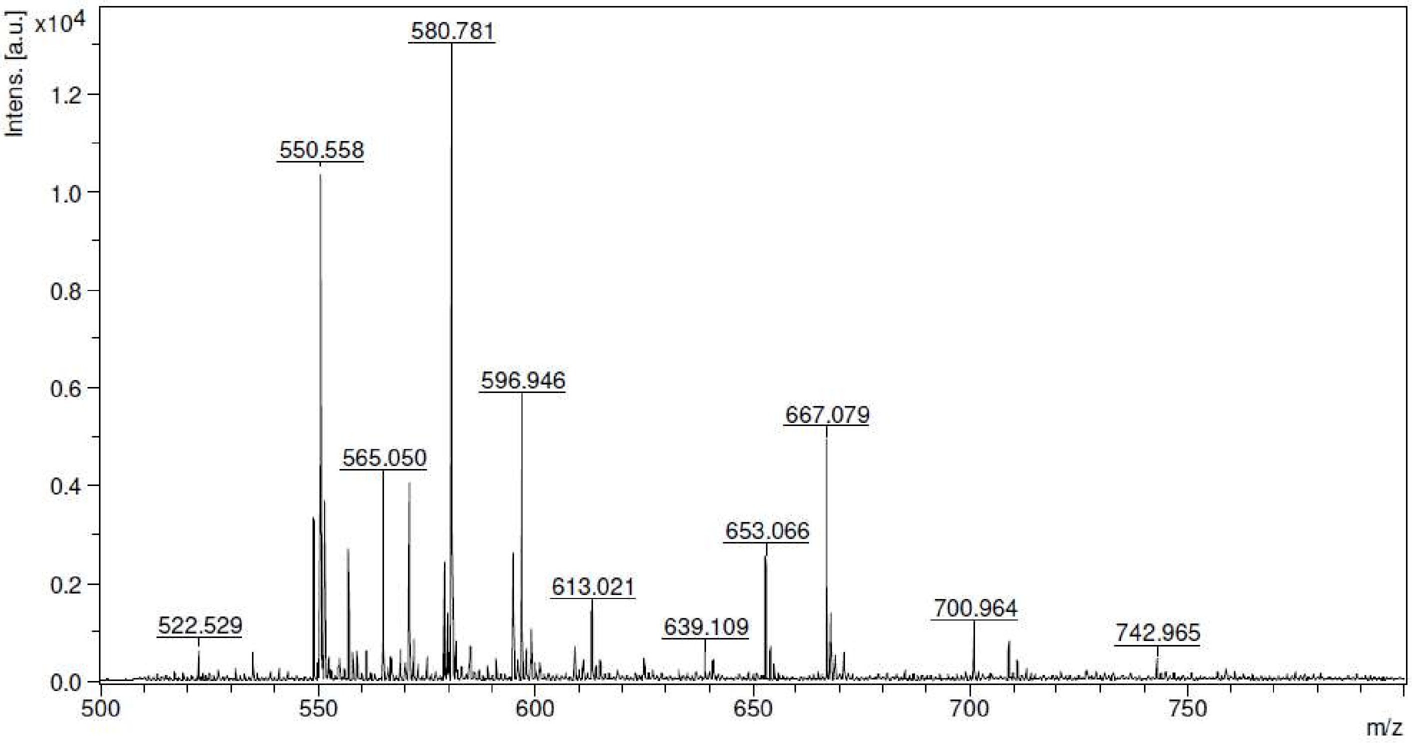


**Figure S1: MALDI-TOF MS spectrum of CJE expanded in m/z range 500-800 amu, positive ion mode, $CsI_2$ added.** For putative assignments see Table S1.

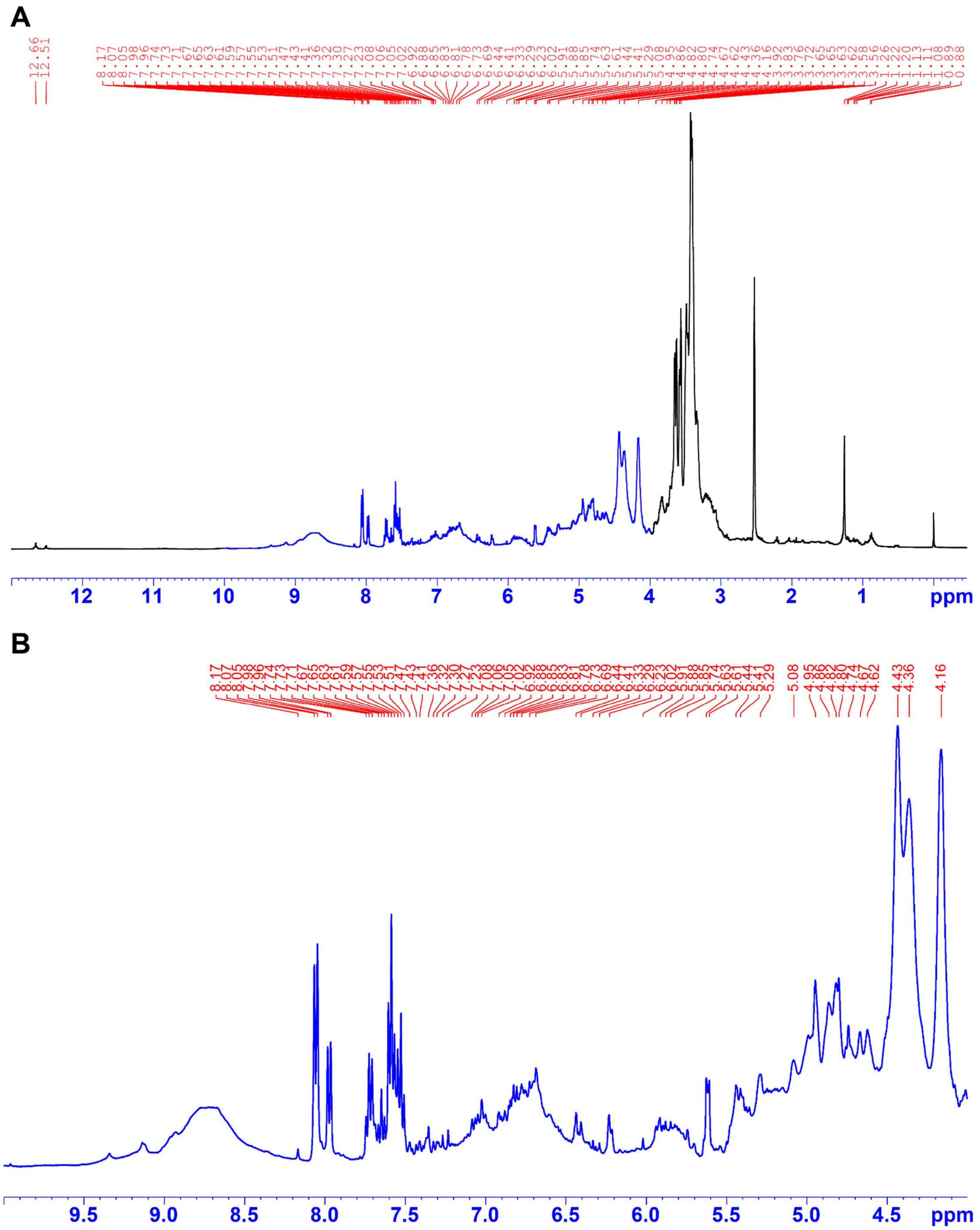


**Figure S2: $^1$H NMR spectrum of CJE in DMSO-$d_6$ full spectrum (A) and expanded in polyphenol signal region (4-10 ppm) (B).**

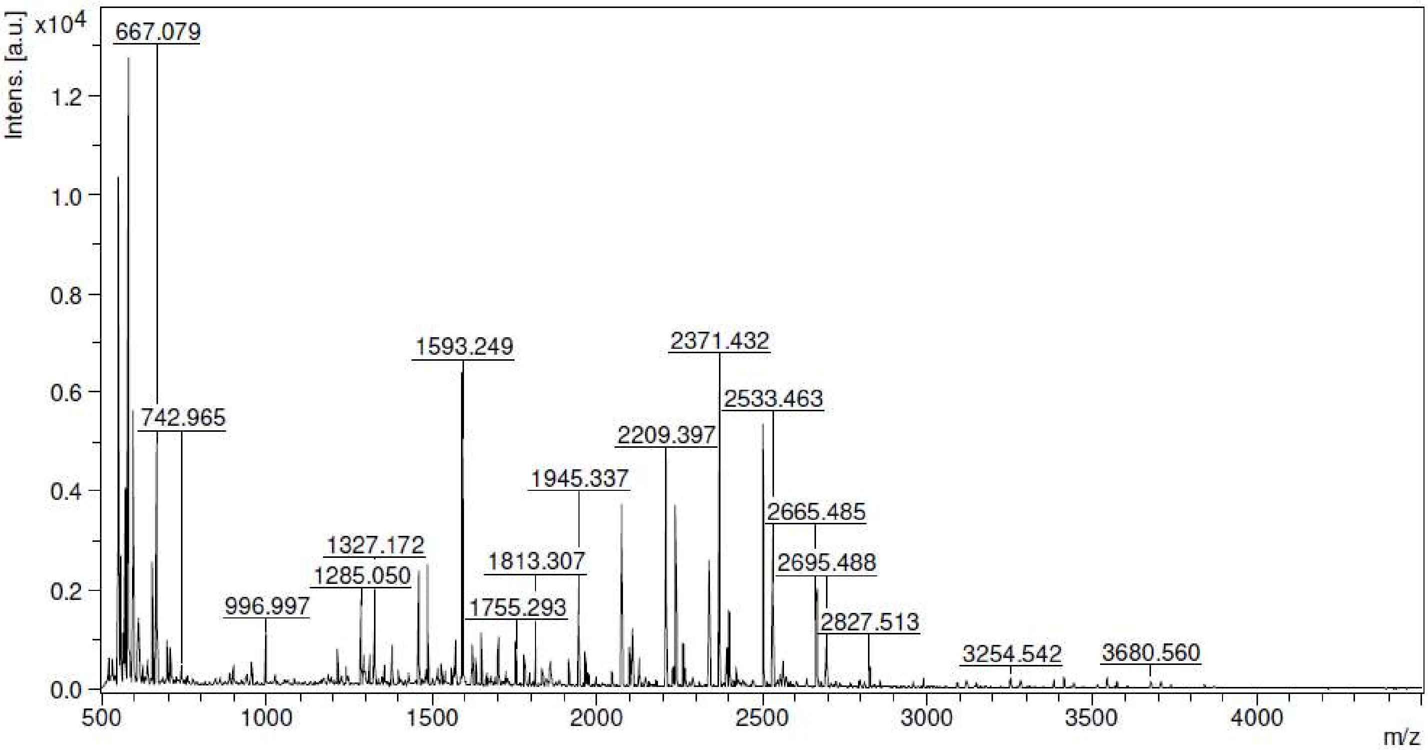


**Figure S3: MALDI-TOF MS spectrum of CJE**. For putative structures see Table S2. Positive ion mode, $CsI_2$ added.

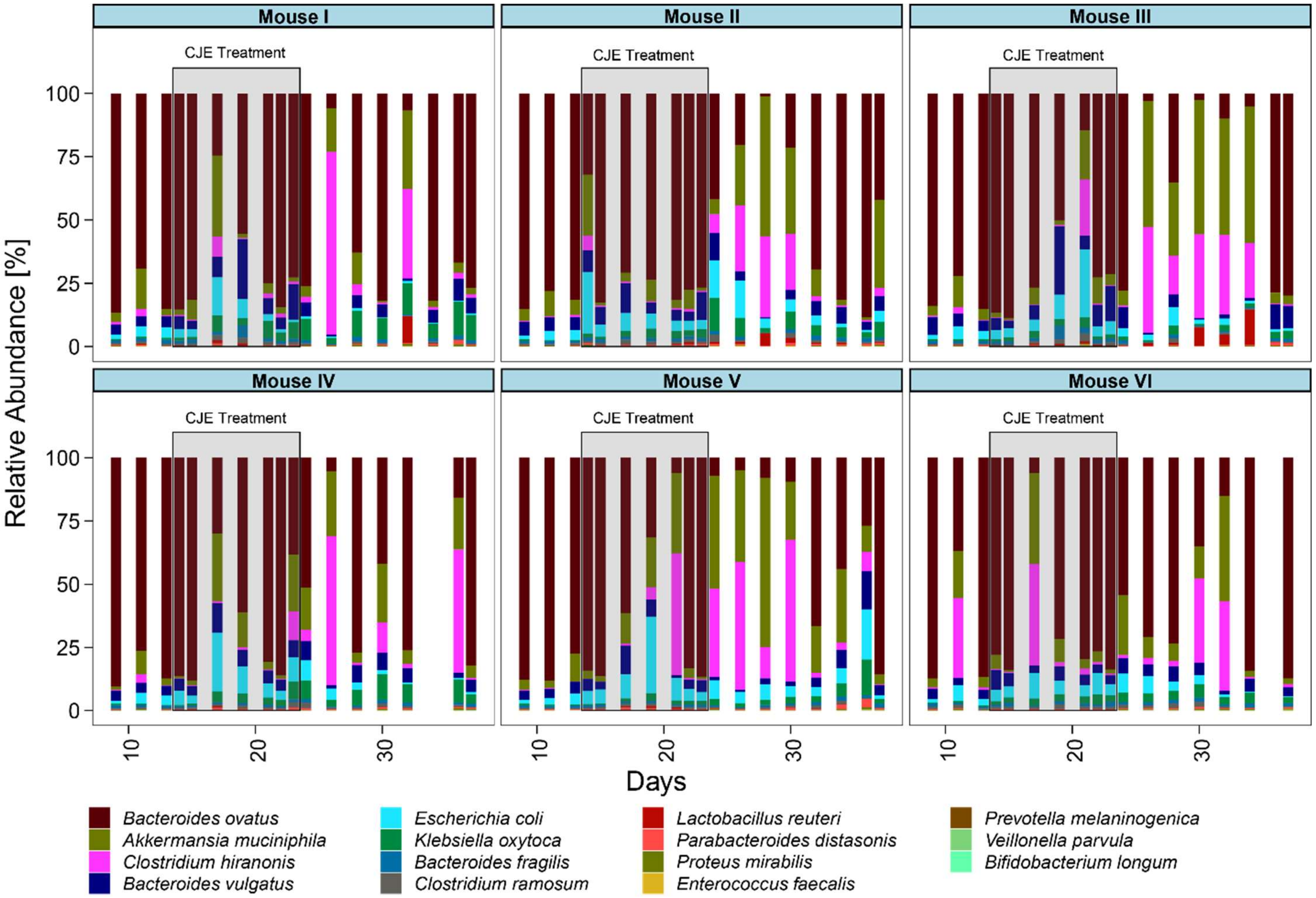


**Figure S4: Relative bacterial abundance for the individual mice throughout the cranberry juice extract (CJE) experiment.**

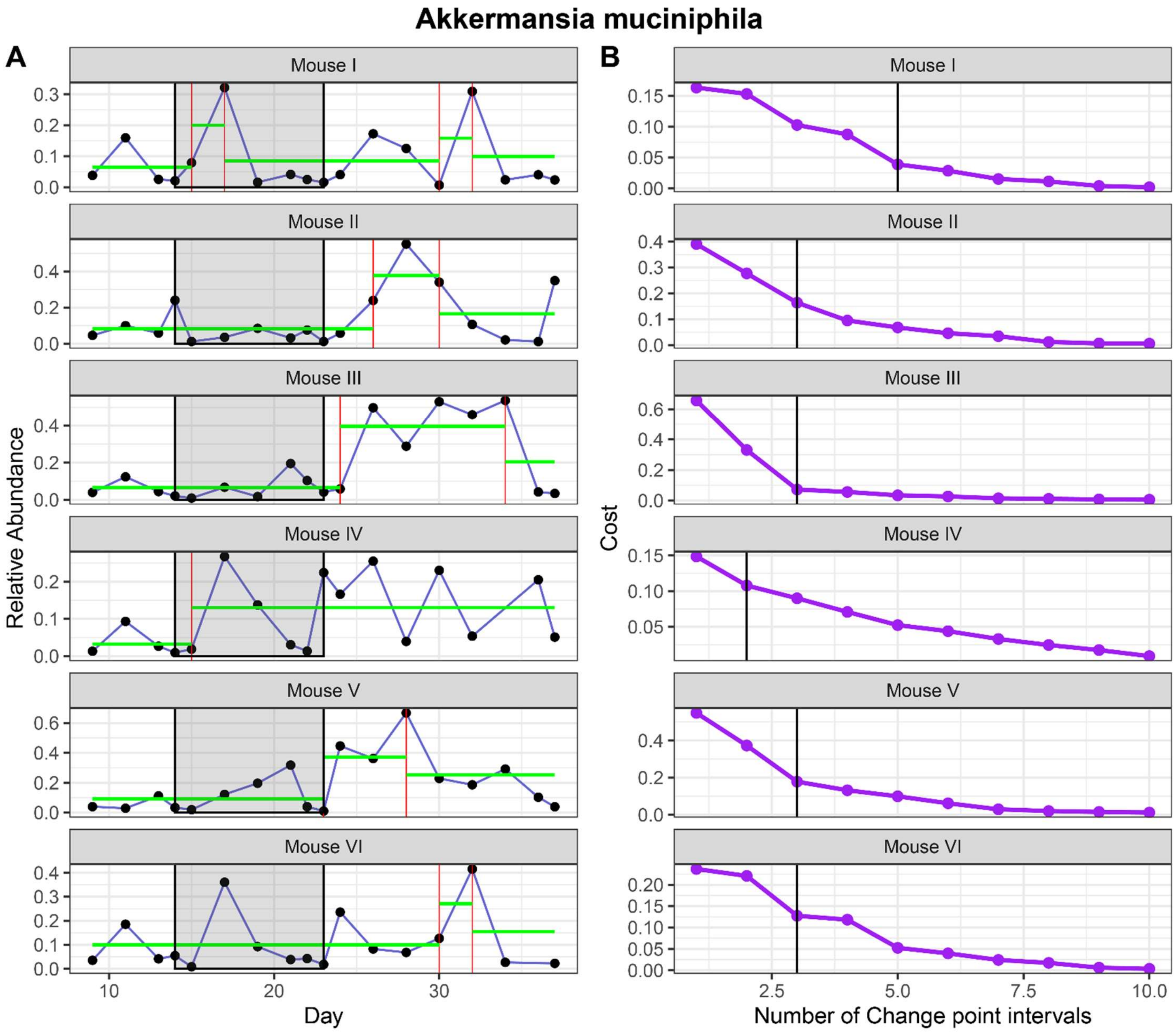


**Figure S5: Change point analysis for *Akkermansia muciniphila* (A) and change point interval determination (B).**

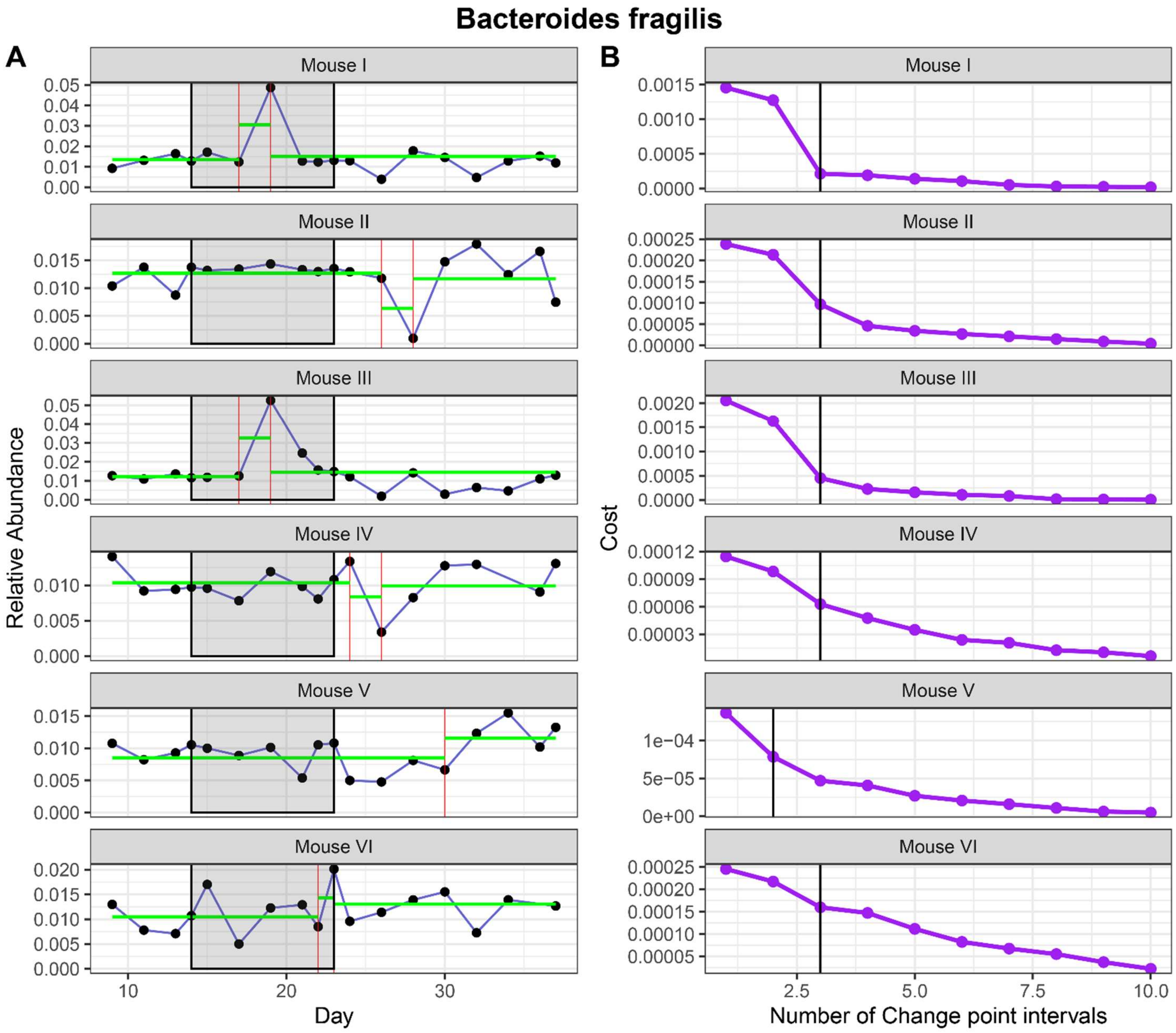


**Figure S6: Change point analysis for *Bacteroides fragilis* (A) and change point interval determination (B).**

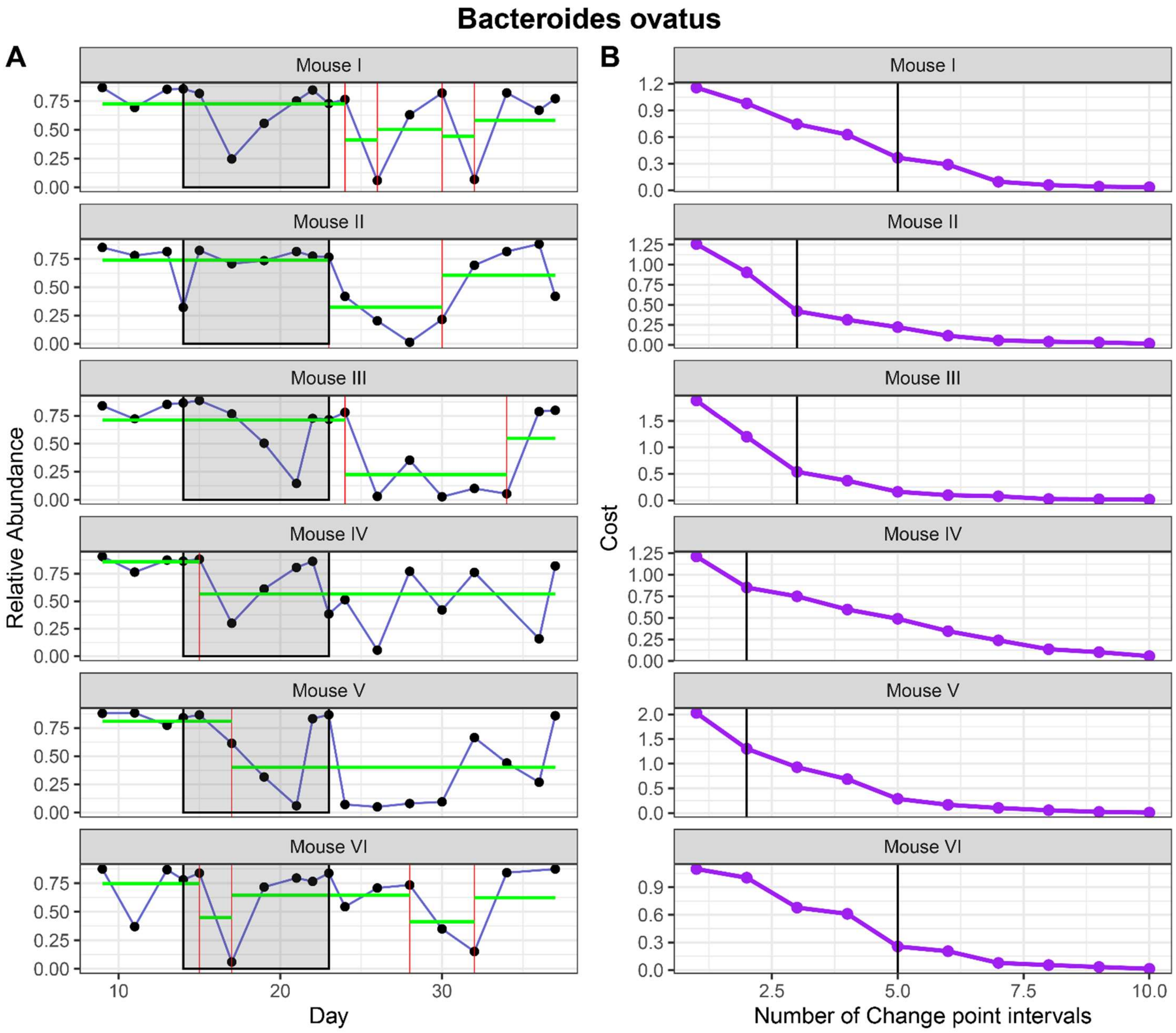


**Figure S7: Change point analysis for *Bacteroides ovatus* (A) and change point interval determination (B).**

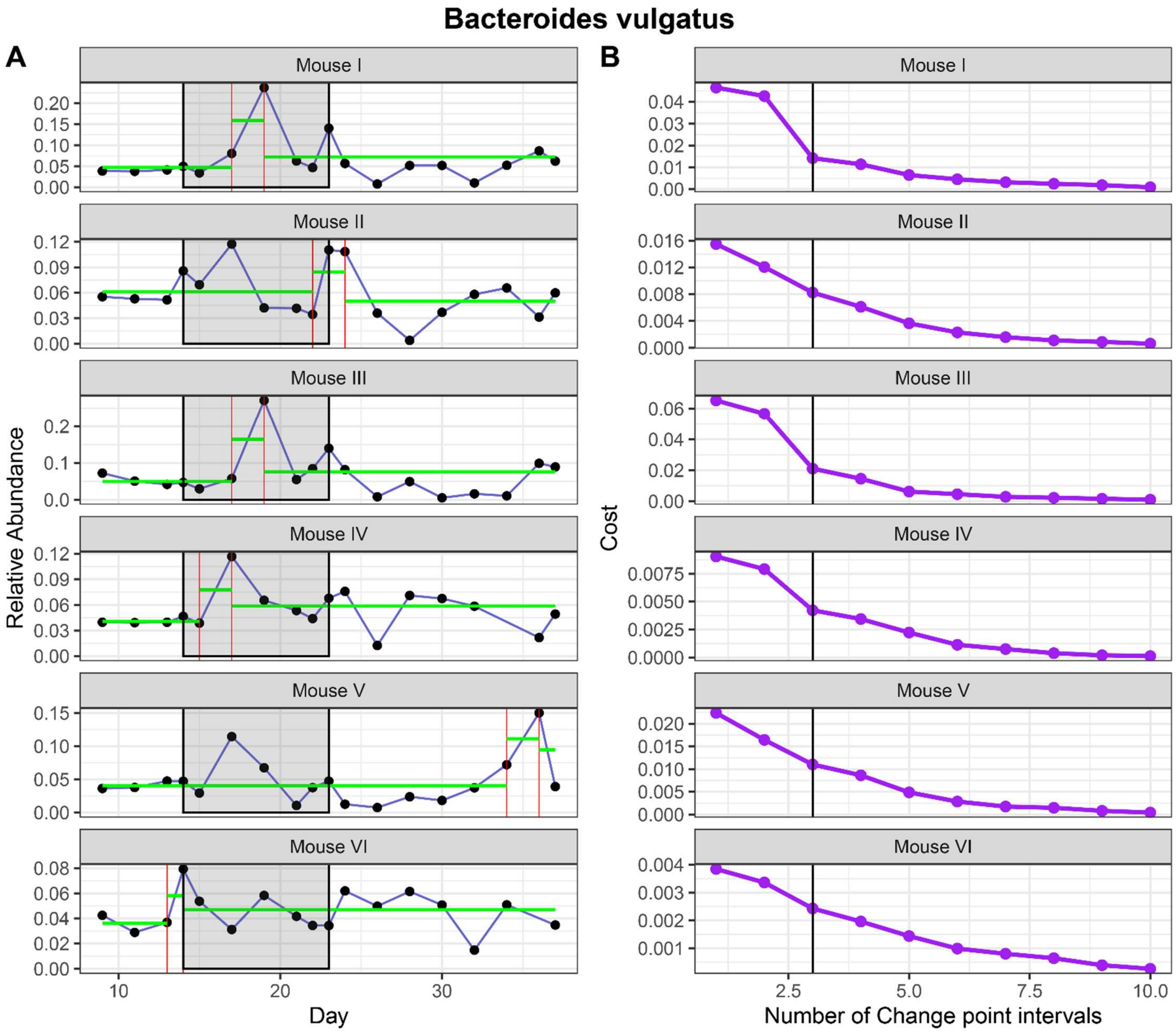


**Figure S8: Change point analysis for *Bacteroides vulgatus* (A) and change point interval determination (B).**

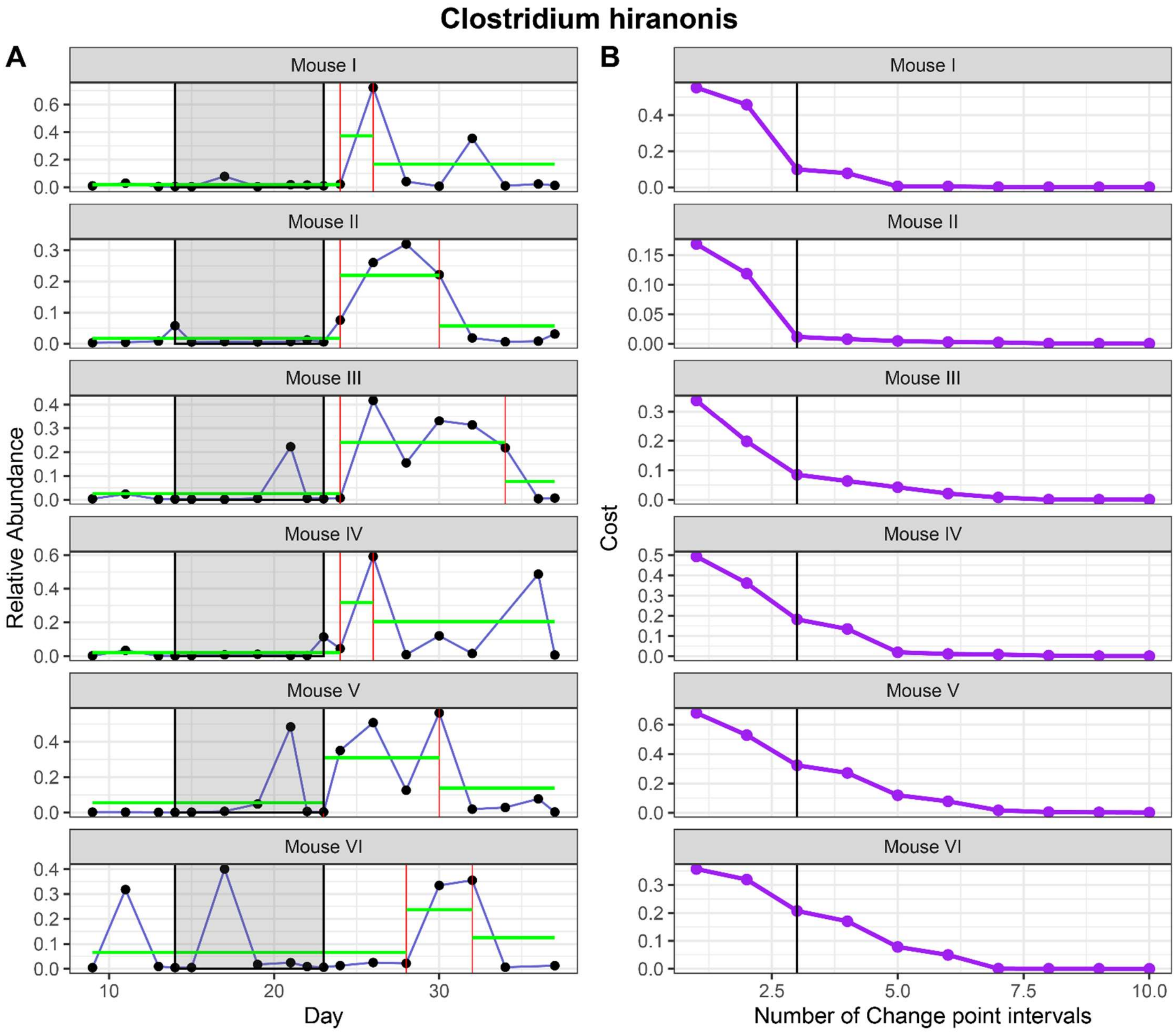


**Figure S9: Change point analysis for *Clostridium hiranonis* (A) and change point interval determination (B).**

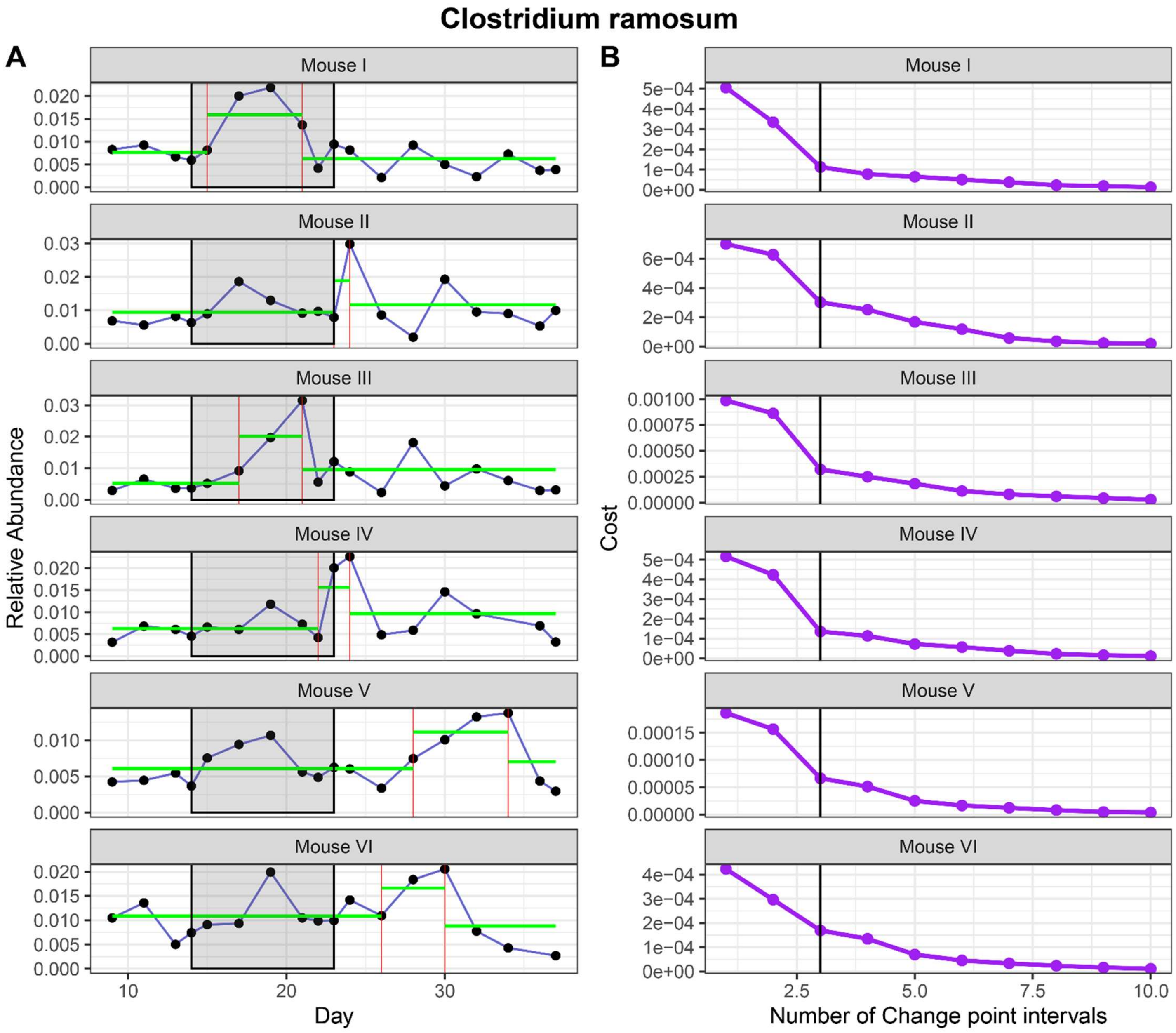


**Figure S10: Change point analysis for *Clostridium ramosum* (A) and change point interval determination (B).**

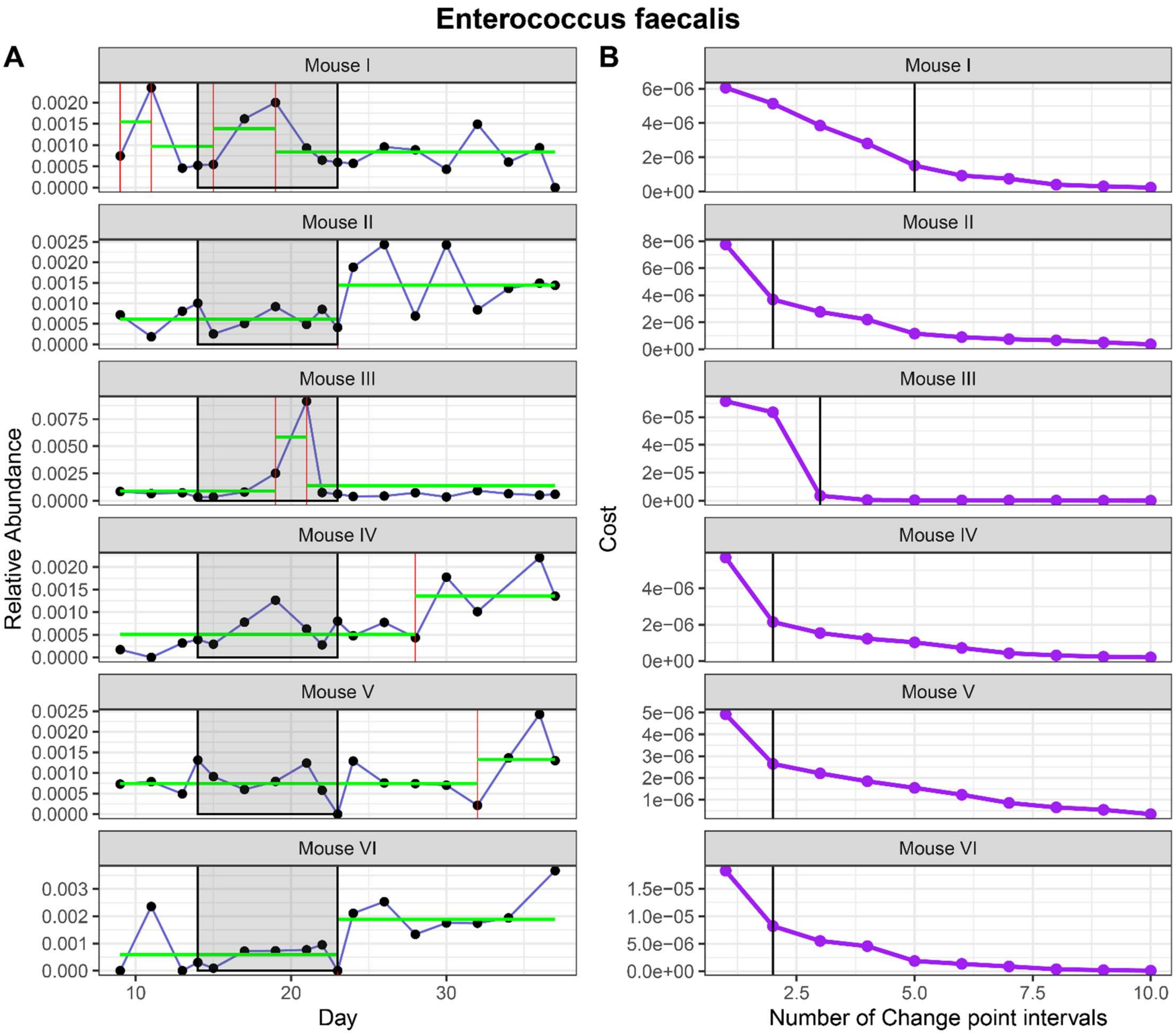


**Figure S11: Change point analysis for *Enterococcus faecalis* (A) and change point interval determination (B).**

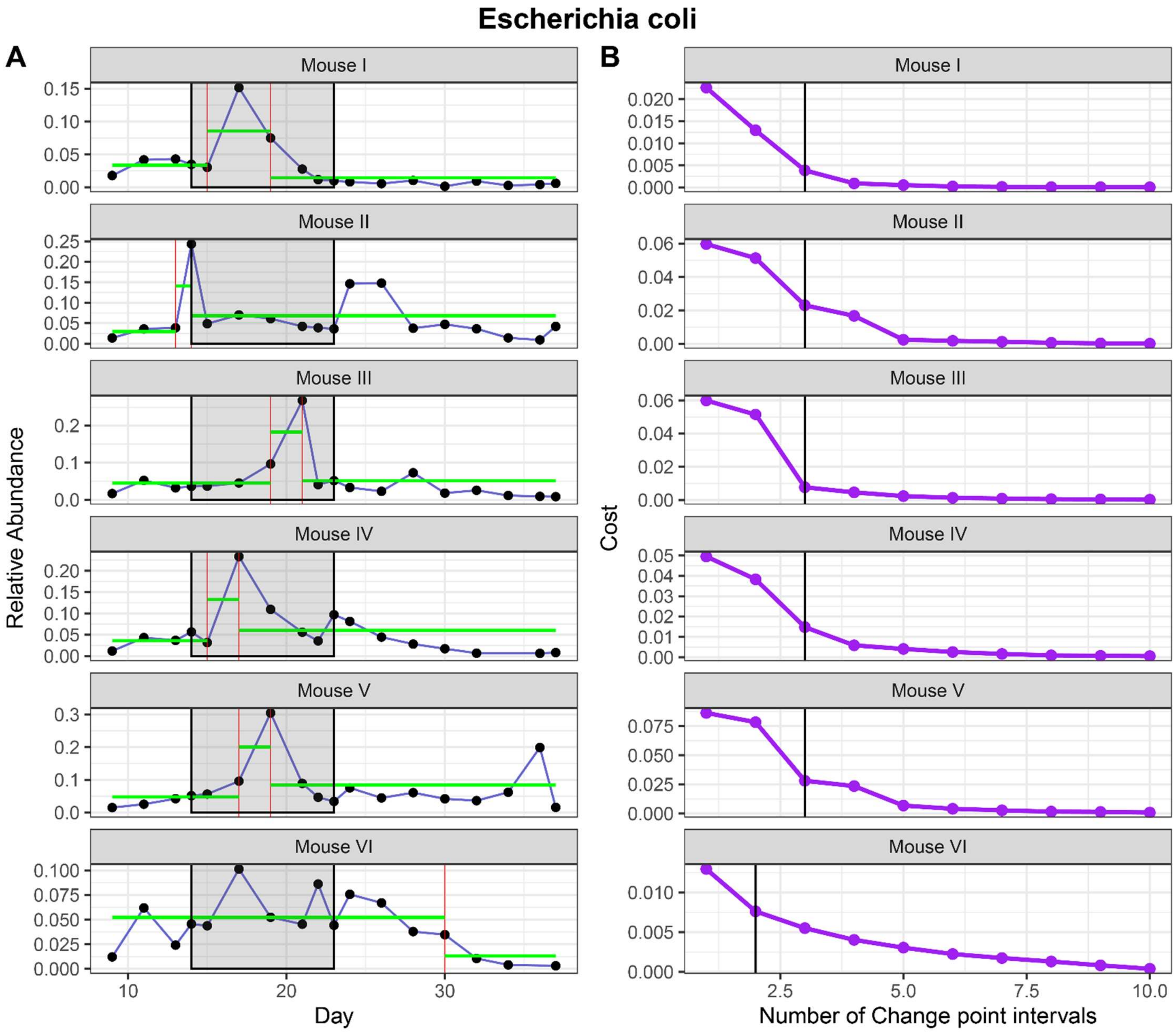


**Figure S12: Change point analysis for *Escherichia coli* (A) and change point interval determination (B).**

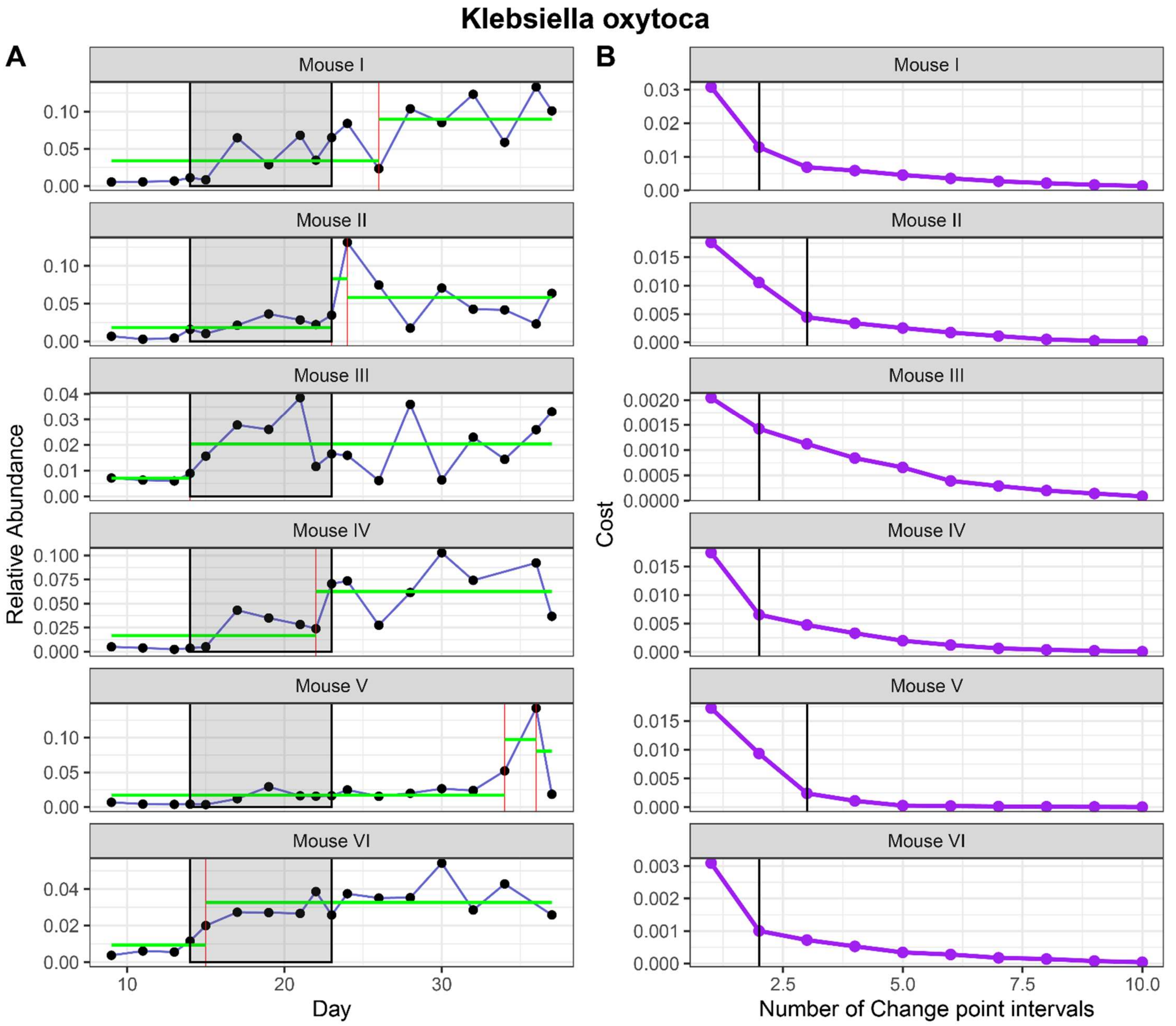


**Figure S13: Change point analysis for *Klebsiella oxytoca* (A) and change point interval determination (B).**

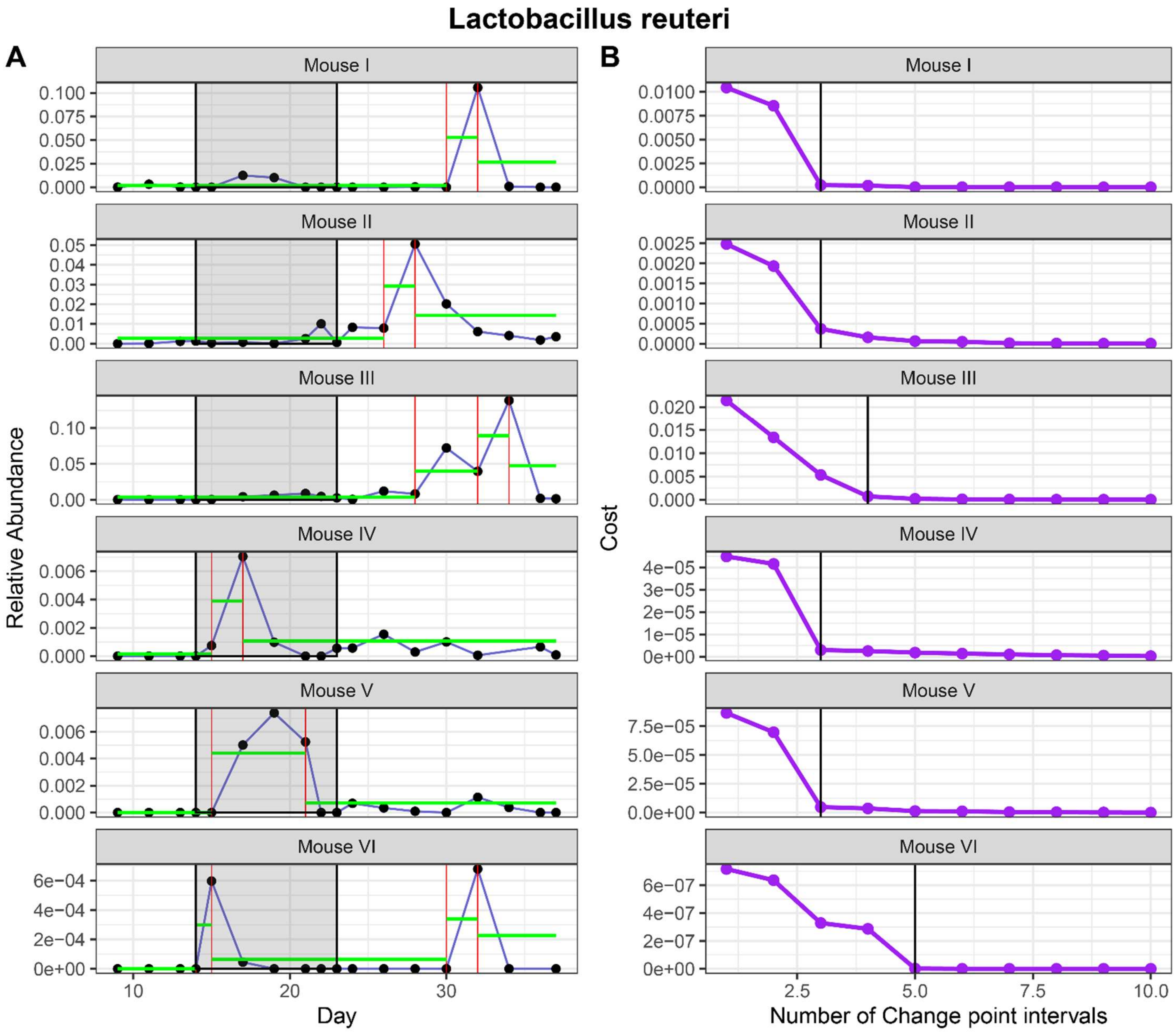


**Figure S14: Change point analysis for *Lactobacillus reuteri* (A) and change point interval determination (B).**

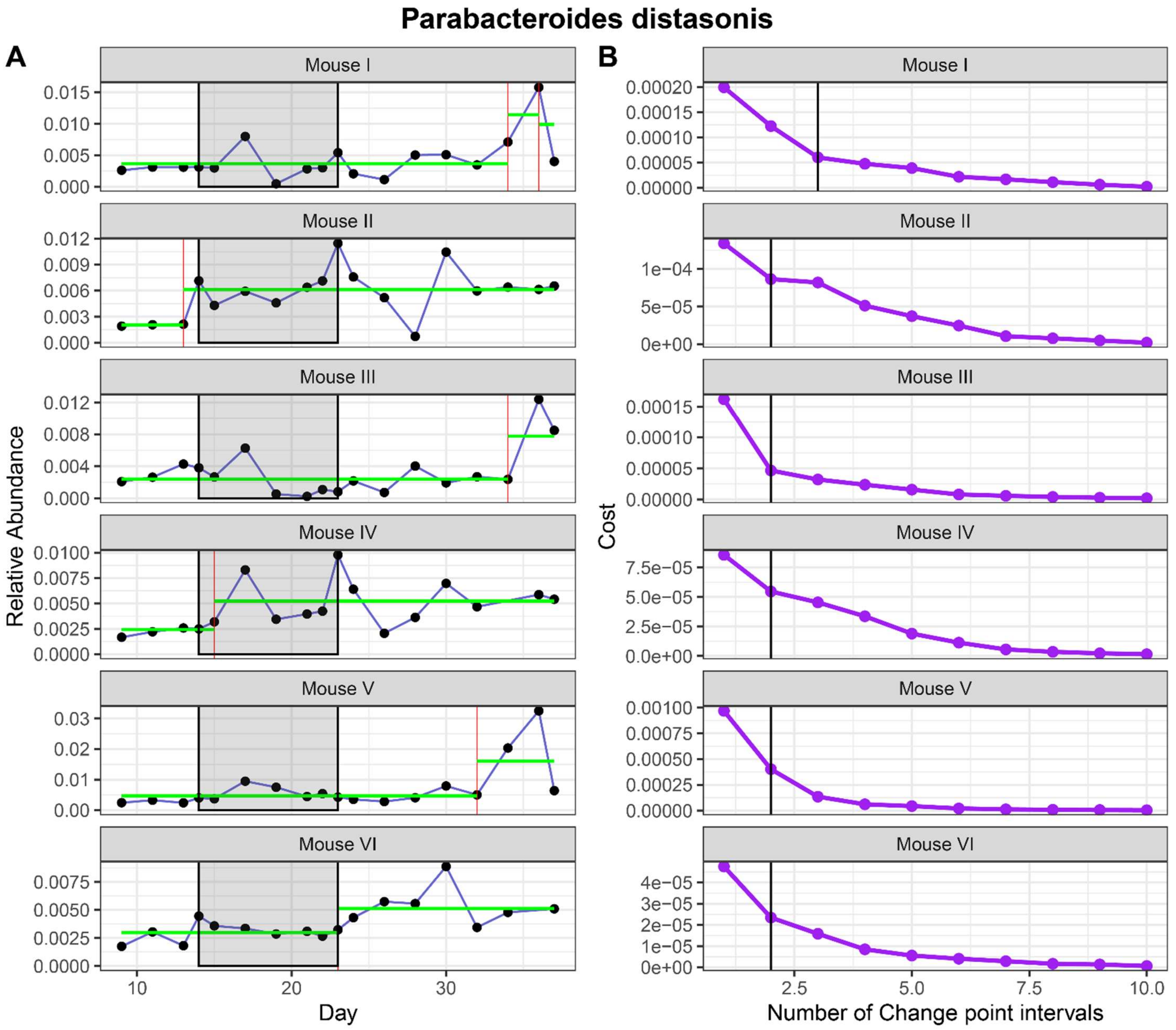


**Figure S15: Change point analysis for *Parabacteroides distasonis* (A) and change point interval determination (B).**

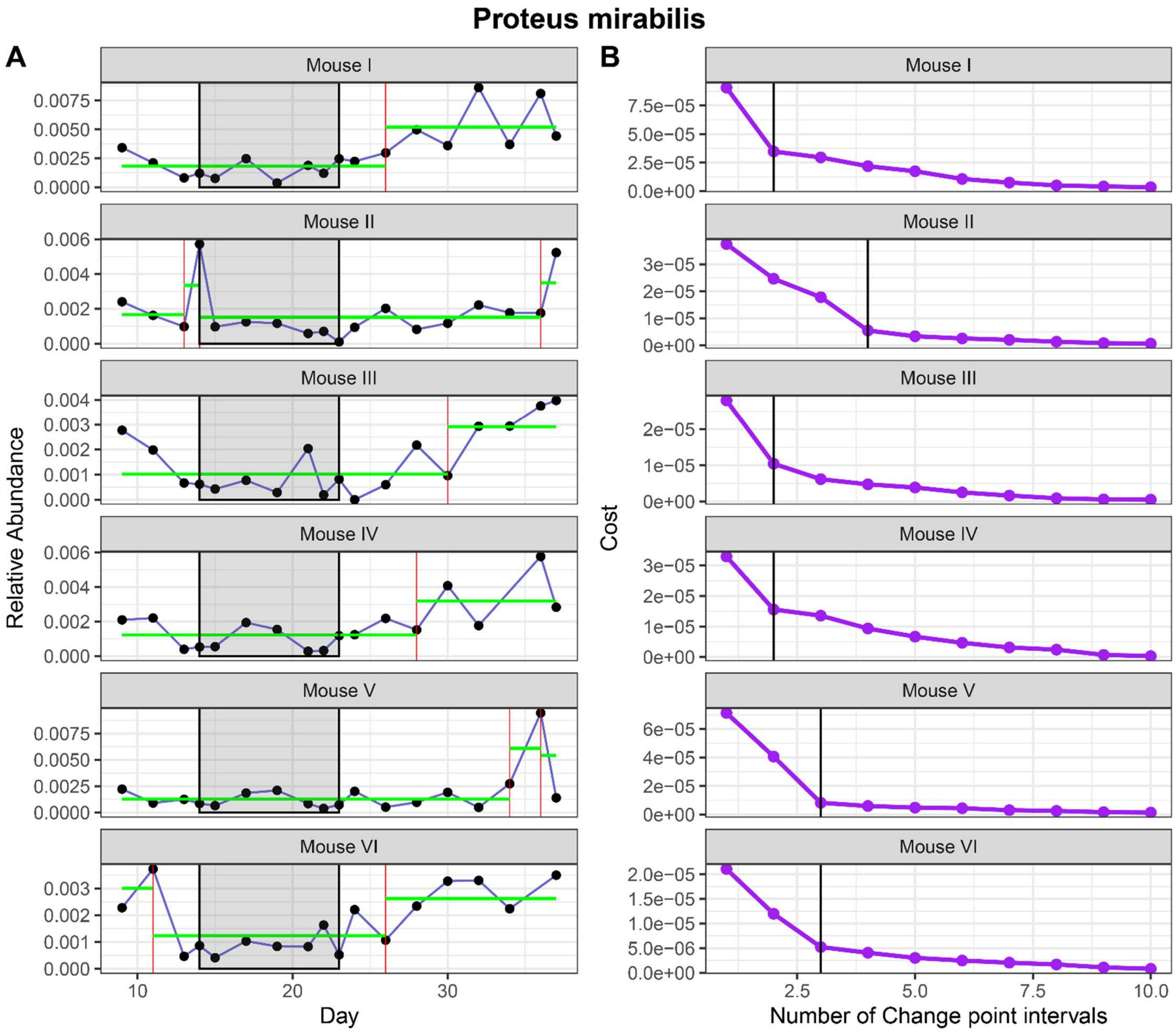


**Figure S16: Change point analysis for *Proteus mirabilis* (A) and change point interval determination (B).**
